# Unique functional responses differentially map onto genetic subtypes of dopamine neurons

**DOI:** 10.1101/2022.12.19.521076

**Authors:** Maite Azcorra, Zachary Gaertner, Connor Davidson, Cooper K. Hayes, Charu Ramakrishnan, Lief Fenno, Yoon Seok Kim, Karl Deisseroth, Richard Longnecker, Rajeshwar Awatramani, Daniel A. Dombeck

## Abstract

Dopamine neurons are characterized by their response to unexpected rewards, but they also fire during movement and aversive stimuli. Dopamine neuron diversity has been observed based on molecular expression profiles; however, whether different functions map onto such genetic subtypes remains unclear. Here, we establish that three genetic dopamine subtypes within the substantia nigra pars compacta each have a unique set of responses to rewards, aversive stimuli, accelerations and decelerations, and these signaling patterns are highly-correlated between somas and axons within subtypes. Remarkably, reward responses were not detected in one subtype, which instead displayed acceleration-correlated signaling. Our findings establish a connection between functional and genetic dopamine subtypes and demonstrate that molecular expression patterns can serve as a common framework to dissect dopaminergic functions.

## Introduction

For decades, midbrain dopamine neurons in the substantia nigra pars compacta (SNc) and ventral tegmental area (VTA) were defined as a largely homogeneous population responding to unexpected rewards and reward-predicting cues^1–7^. However, recent studies have revealed a more complicated story, with increasing evidence for functional heterogeneity. In the VTA, dopamine neurons encode other behavioral variables, such as sensory, motor and cognitive, in addition to the classic reward prediction error response^8^, and separable aversive-responsive populations have been proposed^9,10^. In the SNc, dopamine neurons can respond to both rewarding and aversive stimuli ^11–15^, and increase or decrease firing during movement accelerations ^16–21^.While dopamine neurons and their axons in particular regions of the SNc or striatum respond to these other behavioral variables^16,22^, reward responses have also been observed in the same regions^15,23,24^, leading to the common assumption that most, if not all, dopamine neurons encode reward or reward predicting cues. Therefore, it is currently unclear whether reward, movement and aversion encoding co-occurs in the same neurons or are separately encoded by different groups of dopamine neurons.

Diversity has also been observed in dopamine neurons at the level of gene expression. Previous limitations on the number of molecular markers that could be simultaneously studied resulted in midbrain dopamine neurons being long considered a largely homogeneous population, but recent advances in single cell-transcriptomics have led to the unbiased classification of several putative subtypes ^25–30^. This leads to the enticing hypothesis that different functional responses might in fact map onto different subtypes.

Here we address this question with a focus on the SNc. Three different subtypes have been proposed to account for most of the SNc dopamine neurons^25^, which we here refer to by marker genes that characterizes each subtype: the Aldh1a1+, Calb1+, and Vglut2+ (which is also enriched in Calb1 expression) subtypes. These subtypes have somas in SNc which, while intermingled, are anatomically biased: Aldh1a1+ somas are biased towards ventral SNc, Calb1+ somas towards dorsal SNc, and Vglut2+ somas towards lateral SNc^31^ (Fig. S1A-B, S1E-F). Similarly, their axons project to different regions of striatum, though with overlap in some regions: Aldh1a1+ axons project to dorsal and lateral striatum, Calb1+ to dorso-medial and ventro-medial striatum, and Vglut2+ to posterior striatum^31^ (Fig. S1A, C). If these different subtypes indeed have different functional signaling properties, their anatomical biases might explain previous seemingly conflicting results showing different functional responses of dopamine neurons during the same behaviors^16–24,32^: different subtype(s) may have been inadvertently investigated based on the recording location in SNc or striatum.

## Results

To functionally characterize the different dopamine subtypes, we used intersectional genetic strategies (see Methods) to isolate three known SNc genetic subtypes (Aldh1a1+, Vglut2+, and Calb1+) and label them with the calcium indicator GCaMP6f^33^. We then used fiber photometry to record GCaMP calcium transients from groups of striatal axons of the isolated dopaminergic subtypes in head-fixed mice running on a treadmill while periodically receiving unexpected rewards or aversive stimuli. To control for any movement artifacts, we also recorded GCaMP fluorescence at its isosbestic wavelength, 405 nm^33^. GCaMP is ideally suited for these experiments because all known mechanisms for triggering axonal dopamine release involve increases in intracellular calcium concentration^34,35^, including anterogradely propagating action potentials^36–40^ and cholinergic modulation^41,42^. Critically, the detected calcium transients are generated only from the labeled genetic subtypes; non-labeled neurons do not contribute. For this reason, GCaMP is preferred over extracellular dopamine sensors (i.e. dLight, GRAB-DA, microdialysis) because axons from different subtypes can densely overlap in many striatal regions^31^ and these sensors detect dopamine released from all nearby axons, without subtype specificity.

We will expand and describe the functional signaling properties of the different genetic subtypes in detail in subsequent sections. First, however, we describe a discovery we made about the Aldh1a1+ subtype that prompted us to refine the current genetic classification of dopamine neurons. Given the selective loss of Aldh1a1 neurons in Parkinson’s disease^43,44^, we expected this subtype might show acceleration correlated responses^16^. Functional recordings, however, revealed clear functional heterogeneity across the different recording locations from Aldh1a1+ axons (Fig. S1D-I). Aldh1a1+ axons projecting to dorsal striatum displayed acceleration correlated signaling and no detectable response to rewards (termed a “ Type 1” functional response), while axons projecting more ventrally displaying deceleration correlated signaling and responded robustly to rewards (“ Type 2”) (Fig. S1H-I). This functional heterogeneity was markedly different from recordings from the Calb1+ and Vglut2+ subtypes, which were largely homogenous across recording locations with deceleration correlated signaling and a robust reward response (similar to Aldh1a1+ “ Type 2”). The Type 1 response was remarkable, in that it suggested that there might exist a dopamine neuron subtype that did not respond to rewards, and that showed purely acceleration-correlated responses. If true, this would contradict the notion that all dopamine neurons signal reward. Thus the functional heterogeneity that we observed within Aldh1a1+ recordings motivated us to reexamine the existing dopamine neuron classification schemes and search for new genetic subtypes within the SNc Aldh1a1+ population with such signaling patterns.

### Anxa1+, a new subtype within Aldh1a1+

The current classification of dopamine neurons was derived through single-cell gene expression profiling, primarily via single-cell RNA sequencing (scRNA-seq)^25^. However, such studies are limited by the number of cells analyzed due to technical difficulties in scRNA-seq^45^, which could lead to inconclusive identification of closely related clusters. To uncover more granular divisions among dopaminergic subtypes, we first combined the data from four scRNA-seq studies^27–30^ into an unbiased meta-dataset (see Methods). We observed 8 clusters, one of which was defined by co-expression of Aldh1a1 and Anxa1 (Fig. S2a). Plotting the expression of these two genes showed that Anxa1 expression is limited to a subset of Aldh1a1+ neurons (Fig. S2b). This raised the possibility of at least two molecularly distinct Aldh1a1+ populations. However, while the analysis of this meta-dataset was able to refine our mapping of dopaminergic neuron subtypes, it was still limited by the biases introduced by the individual source datasets and cross-dataset integration methods, and thereby necessitated further validation.

To overcome the technical limitations of single-cell isolation of dopamine neurons, we utilized single-nucleus gene profiling (snRNA-seq), a technique that is more efficient in brain regions where the recovery of intact neurons is difficult^46^. Indeed, this strategy allowed us to profile over 12,000 dopaminergic neuron nuclei from five mice, an order of magnitude higher than previous single-cell studies^26–30^. This approach resulted in the unbiased identification of fifteen clusters, out of which four minor clusters (#12-15 colored in grey in Fig. 1B) represent neurons with weak dopaminergic characteristics (see Methods). The remaining clusters show expression profiles largely in agreement with previous reports from single-cell sequencing studies, but with further subdivision of clusters. Importantly, all clusters were represented in both male and female samples (Fig S3B). Three clusters (#1, #3, #4) were significantly enriched for Sox6 (Wilcoxon Rank Sum test, FDR-adjusted p-values = 4.6 ×10^−150^, 9.8 ×10^−66^, and 2.8 ×10^−276^ respectively); Cluster #2 also showed enrichment of Sox6 (p-value = 1.1 ×10^−04^), however this result did not survive FDR correction. Four clusters (#5, #6, #9, #11) were significantly enriched for Calb1 (FDR-adjusted p-values = 6.6 ×10^−30^, 1.7 ×10^−04^, 1.1 ×10^−22^, 1.5 ×10^−71^, respectively); Cluster 10 also showed Calb1 enrichment (p-value = 6.9 ×10^−04^), which again did not survive FDR correction. Little overlap between these genes was observed (Fig. S3F), recapitulating a fundamental dichotomy among dopaminergic neurons^26,47^. Furthermore, Vglut2 expression is limited to a subset of Calb1+ cells (Fig. 1D, S3I), consistent with prior recombinase-based labelling experiments^31^. We observed two likely SNc clusters with high Aldh1a1 expression (#1 & #4, Fig. 1C, S3I)–the third Aldh1a1+ cluster (#6) was Sox6- and Otx2+ (Fig. 1C, S3F) and corresponds to a previously described VTA subtype also expressing Aldh1a1+^31^. Cluster #4 was again significantly enriched for Anxa1 expression (FDR-adjusted p-value = 9.4 ×10^−118^) (Fig. 1C), corroborating the results from our integrated dataset analysis and establishing Anxa1 as a discrete dopamine neuron subtype marker within Aldh1a1+ neurons.

**Figure 1.**
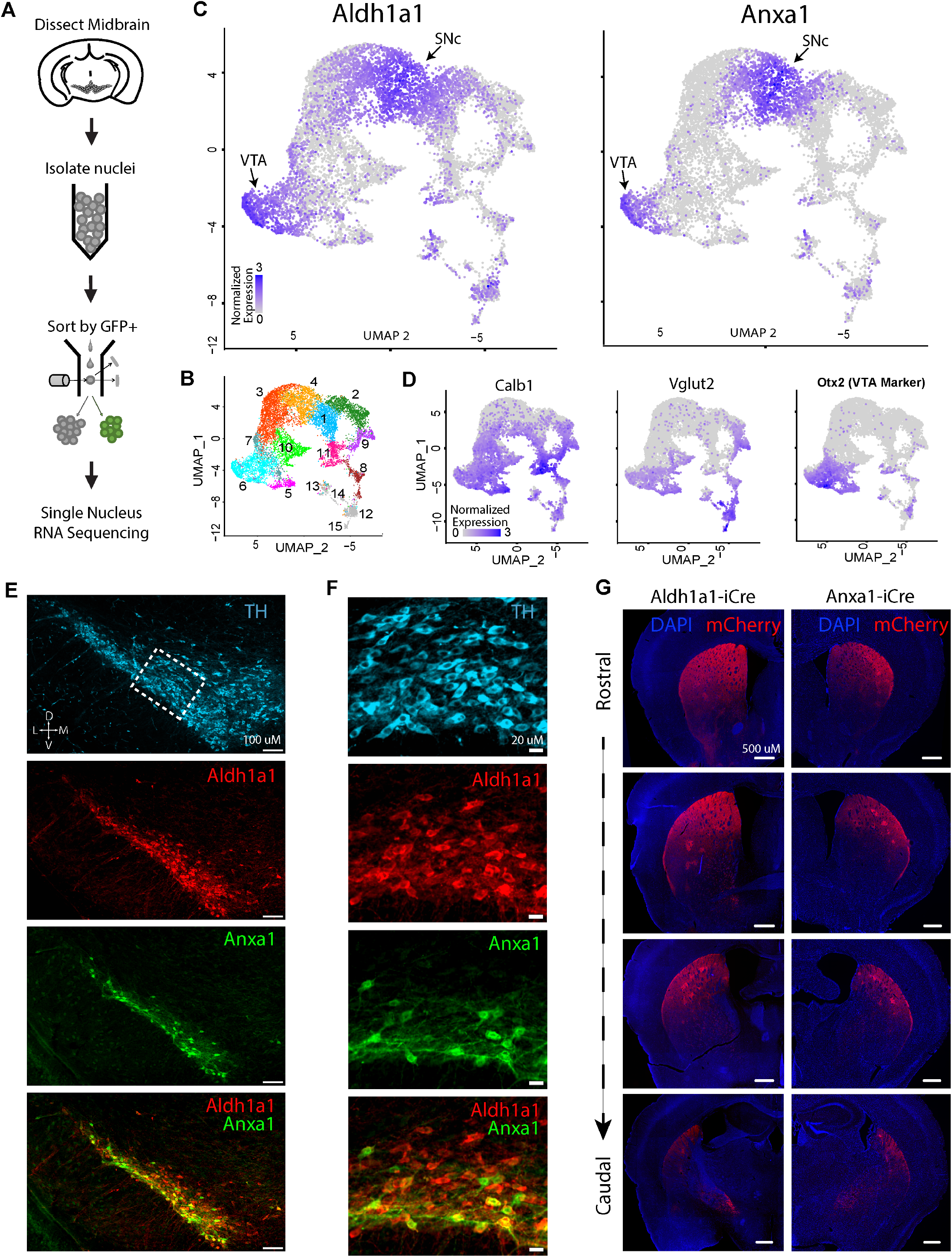
Single-nucleus RNA sequencing reveals an Anxa1-expressing subtype within Aldh1a1+ dopamine neurons. (A) Schematic of snRNAseq experimental pipeline. (B) UMAP reduction of resulting clusters. In total, fifteen clusters were found. Notably, four clusters (12, 13, 14 & 15) had weak dopaminergic characteristics (see Figure S3 for details). (C) Expression of Aldh1a1 and Anxa1, the latter of which is only expressed within a subset of Aldh1a1-expressing neurons. (D) Expression patterns of the additional markers used for genetic access in experiments here, as well as Otx2, a classical marker of most VTA neurons, enriched in clusters 5, 6, and 7. (E) Immunofluorescence images of Aldh1a1 and Anxa1 protein expression in SNc. Anxa1 is limited to a ventral subset of Aldh1a1+ neurons. (F) Zoomed-in crops of section shown in panel E. Anxa1 was ventrally biased within SNc neurons. (G) Right: projection patterns of Anxa1+ SNc axons based on viral labeling, which appear highly-restricted to dorsolateral striatum and patches. Left: projection patterns of Aldh1a1+ SNc axons utilizing the same virus; projections extend more ventrally relative to Anxa1+.

Following the identification of Anxa1+ as a putative subtype, immunostaining confirmed that SNc neurons expressing Anxa1 are indeed part of the broader Aldh1a1+ population, and in fact have cell bodies located ventrally within the already ventral Aldh1a1+ region (Fig. 1E, F). We thus generated a new mouse line, Anxa1-iCre, to genetically access this subtype (see Methods). This allowed us to observe the axonal arbors of Anxa1+ dopamine neurons, which in comparison to Aldh1a1 axon arbors, innervate a more dorsally restricted region of the striatum (Fig. 1G). Importantly, this projection pattern matched the observed anatomical distribution of Aldh1a1+ “ Type 1” axons, suggesting these unique functional responses could map onto the Anxa1+ subtype.

### Genetic subtypes show different signaling patterns during locomotion

To functionally characterize the different dopamine subtypes during locomotion, we used genetic strategies (see Methods) to isolate Vglut2+, Calb1+, and Anxa1+ subtypes (as well as Aldh1a1+ for comparison, and DAT mice where all subtypes were indiscriminately labelled). We then used fiber photometry to record GCaMP calcium transients from populations of striatal axons of isolated dopaminergic subtypes (∼300 micron diameter volumes sampled across the striatal project regions) in head-fixed mice running on a treadmill (Fig. 2A-B). Since the Vglut2+ subtype is contained within Calb1+, in our Calb1+ recordings we avoided recording from the posterior striatum where Vglut2+ neurons project, thus our Calb1+ recordings come largely from Calb1+/ Vglut2-neurons.

**Figure 2.**
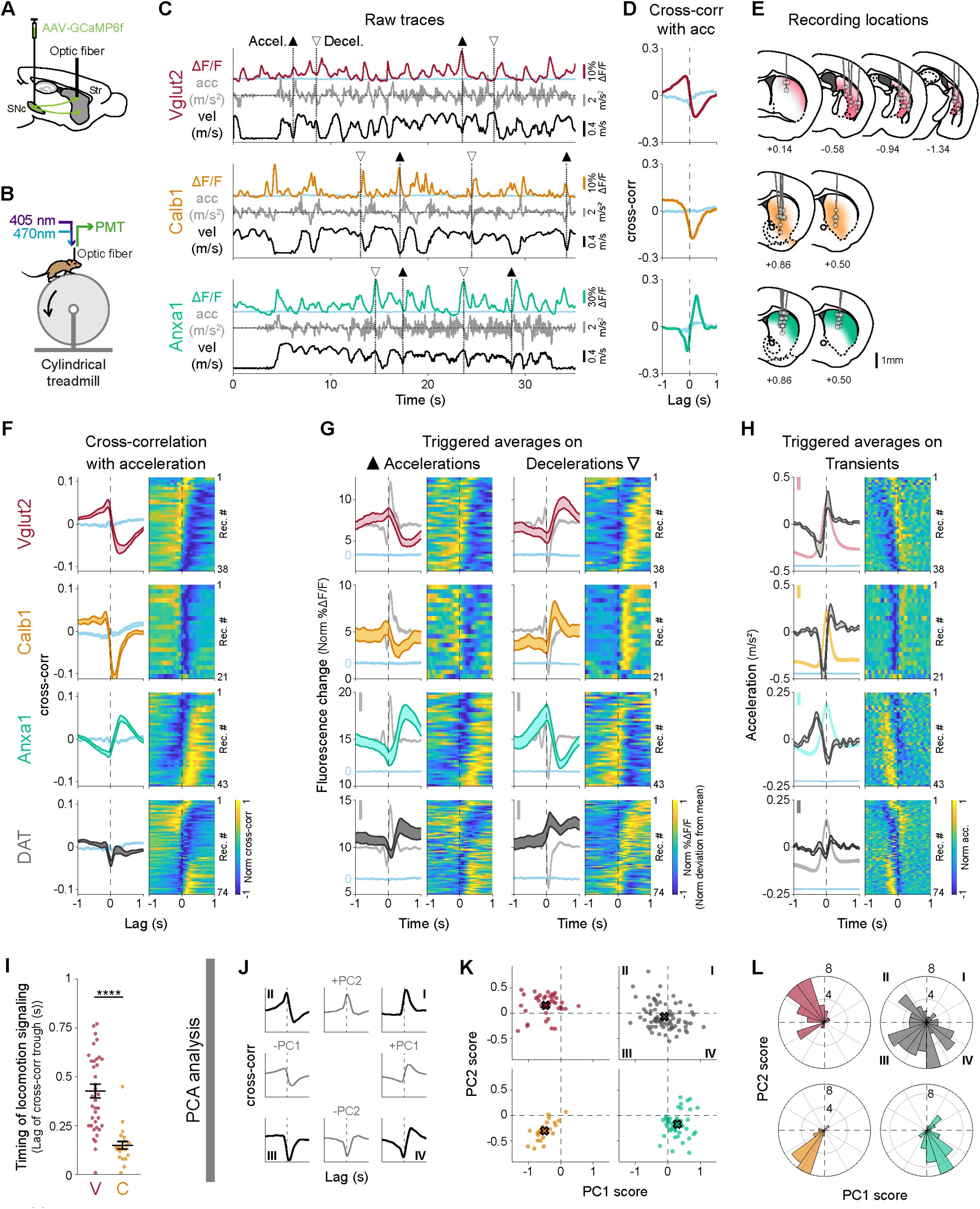
Dopaminergic genetic subtypes display different signaling patterns during locomotion. (A) Strategy used to label dopamine subtypes and record from their axons in striatum with GCaMP6f, a calcium indicator whose changes in fluorescence can be used as a proxy for neuronal firing. (B) Schematic of fiber photometry recording setup. (C) Example recordings from each subtype studied, showing fluorescent traces (ΔF/F), mouse acceleration and velocity. Isosbestic control shown in blue. Large accelerations = ▴, large decelerations = Δ. (D) Cross-correlation between ΔF/F traces and acceleration for traces shown in C. Isosbestic control shown in blue. (E) Recording locations in striatum for recordings shown in F-H. Shaded colors represent projection patterns for each subtype. (F) Average cross-correlation between ΔF/F traces and acceleration for all recordings of each subtype and DAT (subtypes indiscriminately labelled). Isosbestic control shown in blue. Shaded regions denote mean ± s.e.m. Heatmap shows cross-correlations for each recording, sorted by PC1/PC2 angle (see L). Vglut2 mice = 11, n = 38 recordings; Calb1 mice = 5, n = 21; Anxa1 mice = 7, n = 43; DAT mice = 14, n = 74. (G) ΔF/F averages triggered on large accelerations (left, ▴) and large decelerations (right, Δ) for all recordings of each subtype and DAT. Isosbestic control shown in blue, same scale as ΔF/F average but shifted for visibility. Acceleration shown in gray in background (scale bar = 0.2 m/s^2^). Shaded regions denote mean ± s.e.m. Heatmap shows triggered average for each recording, sorted as in F. (H) Acceleration averages triggered on ΔF/F transient peaks for all recordings of each subtype and DAT. ΔF/F average and isosbestic control shown in background (bar = 5% Norm ΔF/F). Shaded regions denote mean ± s.e.m. Heatmap shows triggered average for each recording, sorted as in F. (I) Timing analysis showing the lag of the trough in the ΔF/F-acceleration cross-correlations for each recording from Calb1 and Vglut2. Mean Vglut2 = 0.43, Calb1 = 0.15; p-value for comparison = 6 ×10^−07^ (Wilcoxon rank-sum test with Bonferroni correction). Analogous analysis conducted for triggered averages in Fig. S5B,C. (J-L) Principal component analysis conducted on ΔF/F-acceleration cross-correlations for all striatal recordings from Vglut2, Calb1, and Anxa1 subtypes. (J) Different combinations of PC1 and PC2 loadings representing the different quadrants shown in K-L. Together PC1 and PC2 account for 85% of variance of all cross correlations (PC1 = 63.7% of variance, PC2 = 21.0%). (K) Principal component scores for each recording of each subtype and DAT along PC1 and PC2. X shows mean for each subtype. (L) Radial histogram showing the PC1/PC2 angle of each recording in K. p-values for comparison between subtypes VC = 5 ×10^−07^, VA = 1 ×10^−10^, CA = 3 ×10^−06^ (Wilcoxon rank-sum test with Bonferroni correction).

Remarkably, we observed distinct functional responses in DA neuron subtypes. Calb1+ and Vglut2+ axons preferentially signaled during locomotion decelerations, while Anxa1+ axons preferentially signaled during locomotion accelerations (Fig. 2C), similarly to Aldh1a1+ “ Type 1”. Accordingly, cross-correlations between calcium ΔF/F traces (ΔF/F traces) and acceleration revealed a deep trough at positive time lags for Calb1+ and Vglut2+ axons (indicative of calcium transients following decelerations), but a large peak at positive lags for Anxa1+ axons (transients following accelerations) (Fig. 2D), and this was consistent across a wide range of striatum locations (Fig. 2E-F). The opposing signaling patterns of Calb1+ and Vglut2+ vs Anxa1+ was also clear in ΔF/F averages triggered on accelerations or decelerations (Fig. 2G) and acceleration averages triggered on ΔF/F transient peaks (Fig. 2H). Importantly, these signaling differences persist even in regions where axons from different subtypes overlap (Fig. 3A), indicating that these functional differences were intrinsic to each subtype, and were not simply defined by striatal projection location. In contrast, in DAT mice where subtypes were indiscriminately labelled, heterogeneous signaling was observed across striatal recording locations (Fig. 2F-H bottom, S5A; and to a lesser extent in Aldh1a1+, Fig. S5D-F).

**Figure 3.**
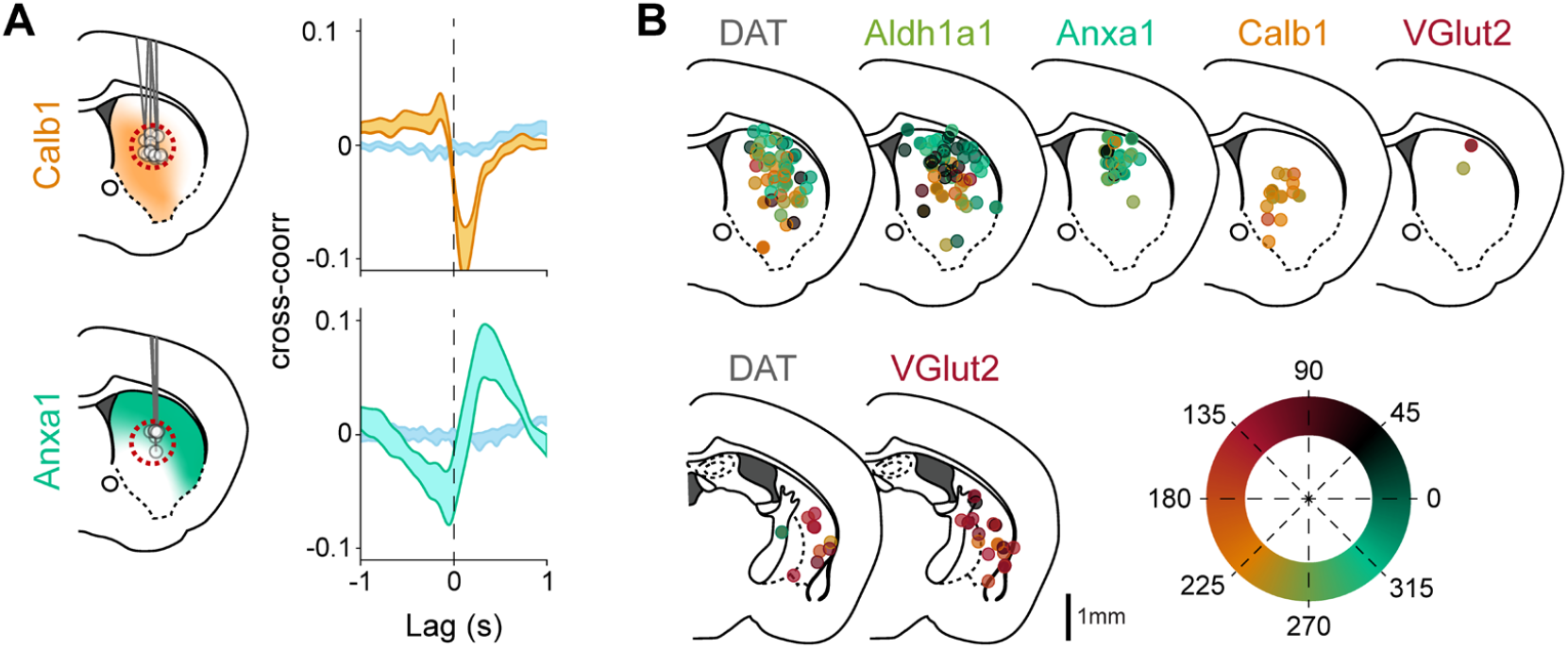
Spatial distribution of subtype specific locomotion responses. (A) Comparison of locomotion response (cross-correlation between ΔF/F and acceleration) for Calb1 and Anxa1 recordings only from a region of striatum where their axons overlap, dashed red circle. Isosbestic controls in blue. Shaded regions denote mean ± s.e.m. Calb1 mice = 4, n = 12 recordings, Anxa1 mice = 4, n = 7. (B) Locomotion response (PC1/PC2 angle, as shown in Fig. 2L) mapped onto recording location for each subtype and DAT. Locations from the body (top) or the tail of the striatum (bottom) were collapsed into a single brain section. To reduce overlap, locations were shifted a random amount between ±0.4mm mediolaterally.

Interestingly, signaling differences were also evident between Vglut2+ and Calb1+ in their timing with respect to decelerations, with Calb1+ transients following decelerations with a shorter lag than Vglut2+ (Fig. 2I, Fig. S5B-C). To further quantify such differences, we used a dimensionality reduction technique to extract the components that best explain the variance in the cross-correlations. We applied principal component analysis (PCA) to the matrix of all cross-correlation traces from Vglut2+, Calb1+, and Anxa1+ subtypes (see Methods), finding that the first two principal components (PC1 and PC2) explained 85% of the variance in the cross-correlations (64% PC1 and 21% PC2). We observed that different combinations of PC1 and PC2 closely approximated the cross-correlation averages of the different subtypes: PC1^+^ + PC2^-^ for Anxa1+, PC1^-^ + PC2^-^ for Calb1+, and PC1^-^ + PC2^+^ for Vglut2+ (Fig. 2J, H). Accordingly, the decomposition of each recording along these principal components revealed distributions that were well separated between the subtypes (Fig. 2K-L. Mean PC1/PC2 angles, representing the time-course of the cross-correlations and thus the temporal relationship between ΔF/F and acceleration = 141° for Vglut2+, 218° for Calb1+, 244° for Anxa1+, p-values = 5 ×10^−07^ V-C, 1 ×10^−10^ V-A, 3 ×10^−06^ C-A, Wilcoxon rank-sum test with Bonferroni correction). Cross-correlations from DAT recordings decomposed using the same principal components were spread across the same regions of the PC1/PC2 space as individual subtypes, and areas in between (Fig. 2K-L, dark grey; and to a lesser extent in Aldh1a1+, Fig. SG-H). These different decompositions of DAT (and Aldh1a1+) recordings also mapped onto different striatal locations (Fig. 3B). DAT recordings that displayed similar decomposition to a particular subtype (e.g. dorsal striatum to Anxa1+ or posterior striatum to Vglut2+) suggest that a single subtype dominated DAT signaling within the photometry recording volume in these striatal regions. However, the DAT recordings that displayed a different mixture of PCs than any particular subtype (e.g. middle depth striatum) suggest that a mixture of subtype axons were contained within the recording volume (Fig. 3B). Overall, these findings demonstrate that during locomotion Calb1+, Vglut2+ and Anxa1+ dopamine neuron subtype axons displayed different average functional signaling patterns; Calb1+ and Vglut2+ axons were largely deceleration correlated with unique timing differences between these subtypes, while Anxa1+ axons were largely acceleration correlated.

### Subtypes show different responses to rewards and aversive stimuli

We then asked whether these dopaminergic subtypes respond differently to rewards and aversive stimuli. We randomly delivered unexpected water rewards and aversive air puffs to the whiskers/face to mice already habituated to run on the treadmill (Fig. 4A), and again used fiber photometry to record ΔF/F transients from populations of axons at different striatal locations (Fig. 4B). We found that both Calb1+ and Vglut2+ axons responded robustly to rewards (Fig. 4C-E, G; 0.5s cumulative response: mean = 7.9 and 13.4 Norm %ΔF/Fs, p-values = 2 ×10^−04^ and 2 ×10^−04^, respectively, Wilcoxon Signed Rank test with Bonferroni correction) and air puffs (Fig. 4C, F-G; 0.5s cumulative response: mean = 15.3 and 6.4, p-values = 1 ×10^−04^ and 2 ×10^−04^, respectively) consistently across nearly all recording locations. The reward signaling in Calb1+ and Vglut2+ axons could not be explained by their responses to mouse movement during reward delivery, as similar results were obtained for reward delivery during rest (Fig. S6A-B). For air puffs however, it was not possible to exclude this possibility since mice invariably moved in response to the aversive stimulus.

**Figure 4.**
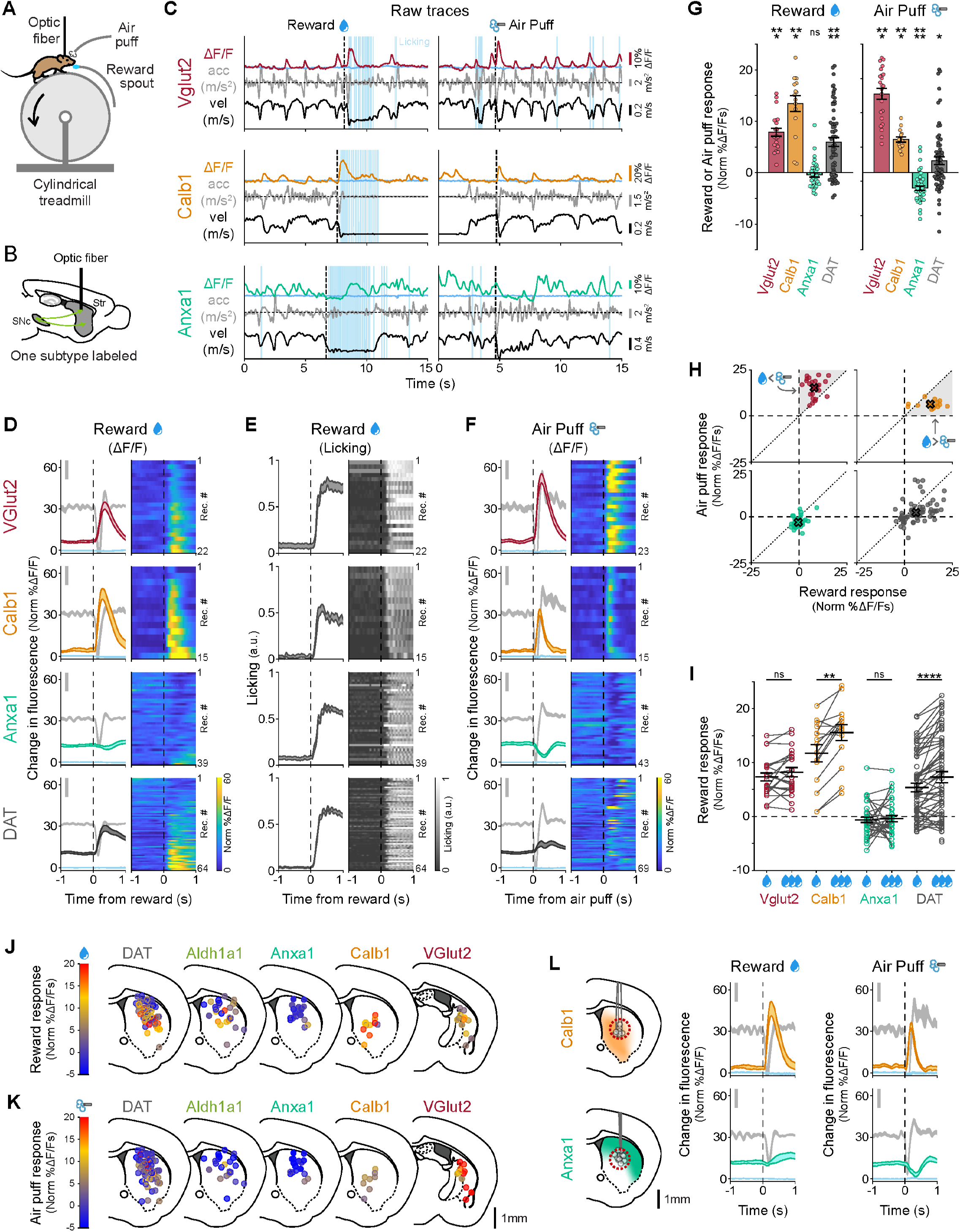
Dopaminergic genetic subtypes display different responses to rewards and aversive stimuli. (A) Mouse running on treadmill during fiber photometry while receiving unexpected rewards and air puffs. (B) Schematic of fiber photometry recording strategy. (C) Example recordings for each subtype studied, showing fluorescence traces (ΔF/F), mouse velocity, acceleration, licking, and reward (left) or air puff (right) delivery times. Isosbestic controls in light blue, same scale as ΔF/F traces. Reward and Air puff examples for each subtype are from the same recording. (D) ΔF/F averages triggered on reward delivery times for all recordings of each subtype and DAT. Isosbestic control in light blue, same scale as ΔF/F average. Acceleration shown in gray in background (scale bar = 0.2 m/s^2^). Shaded regions denote mean ± s.e.m. Heatmaps show triggered average for each recording, sorted by size of reward response. Vglut2 mice = 10, n = 22 recordings; Calb1 mice = 7, n = 15; Anxa1 mice = 6, n = 39; DAT mice = 12, n = 64. (E) Licking average triggered on reward delivery times for all recordings of each subtype and DAT (same as D). Shaded regions denote mean ± s.e.m. Heatmap shows triggered average for each recording, sorted as in D. (F) ΔF/F averages triggered on air puff delivery times for all recordings of each subtype and DAT. Isosbestic control in light blue, same scale as ΔF/F average. Acceleration shown in gray in background (scale bar = 0.2 m/s^2^). Shaded regions denote mean ± s.e.m. Heatmap shows triggered average for each recording, sorted by reward size as in D,E. Vglut2 mice = 11, n = 23 recordings; Calb1 mice = 7, n = 15; Anxa1 mice = 7, n = 43; DAT mice = 12, n = 69. (G) Average reward and air puff responses for each subtype (integral of fluorescence in a 0.5 s window after stimulus minus integral in 0.5 s before stimulus). Error bars denote ± s.e.m. Means and p-values for reward: Vglut2 m = 7.9 Norm ΔF/Fs, p = 2 ×10^−04^; Calb1 m = 13.4, p = 2 ×10^−04^; Anxa1 m = -0.4, p = 0.3 (not significant); DAT m = 6.0, p = 5 ×10^−07^. Means and p-values for air puff: Vglut2 m = 15.3, p = 1 ×10^−04^; Calb1 m = 6.4, p = 2 ×10^−04^, Anxa1 m = -3.0, p = 2 ×10^−05^; DAT m = 2.3, p = 0.02. Wilcoxon signed-rank test with Bonferroni correction. (H) Reward vs air puff responses for all recordings of each subtype and DAT. X shows mean for each subtype. Shaded regions are areas representing greater air puff than reward response (for Vglut2) or greater reward vs air puff response (for Calb1). (I) Comparison of responses to small vs large rewards for each subtype. Error bars denote mean ± s.e.m. Mean difference and p-values: Vglut2 m = 0.9 Norm ΔF/Fs, p = 0.2 (not significant); Calb1 m = 3.8, p = 2 ×10^−04^; Anxa1 m = 0.2, p = 1 (not significant); DAT m = 1.9, p = 7 ×10^−06^. Paired Wilcoxon Signed Rank test with Bonferroni correction. (J) Reward response mapped onto recording locations for each subtype and DAT. Locations from the body or the tail of the striatum were collapsed into a single brain section. To reduce overlap, locations were shifted a random amount between ±0.4mm mediolaterally. (K) Same as J but for air puff response. (L) Comparison of reward and air puff response for Calb1 and Anxa1 recordings only from a region of striatum where their axons overlap, dashed red circle. Isosbestic control in blue. Shaded regions denote mean ± s.e.m. Calb1 mice = 3, n = 8 recordings, Anxa1 mice = 5, n = 15.

In contrast, however, unexpected reward responses were not detectable from Anxa1+ axons (Fig. 4C-E, G; 0.5s cumulative response: mean = -0.4, p-value = 0.3, not significant), but they did respond to air puffs with a signaling decrease (Fig. 4C, F-G; 0.5s cumulative response: mean = -3.0, p-value = 2 ×10^−05^), though again this response could have been due to mouse movement. Importantly, these differences persist even in regions where axons from different subtypes overlap (Fig. 4L), indicating that they were intrinsic to each subtype and not simply defined by striatal projection location. 3 of 39 Anxa1+ recordings locations displayed a small increase in ΔF/F post-reward (Fig. 4D, Anxa1+, bottom rows), however these were likely movement responses since no increases were observed when rewards were delivered at rest (Fig. S6A-B). The reward and air puff responses from DAT recordings were location specific, with few reward responses in dorsal striatum, but responses more prevalent in more ventral and posterior regions (Fig. 4J-K).

Interestingly, while the Vglut2+ and Calb1+ subtype axons both responded to rewards and air puffs, their responses still differed. Vglut2+ axons displayed larger responses to air puffs than reward while Calb1+ axons displayed larger responses to rewards than air puffs (Fig. 4H). Furthermore, Calb1+ axons displayed larger responses to increased reward size–a hallmark of reward prediction error (RPE)^48,49^. This response increase was not detectable from Vglut2+ axons (Fig. 4I). Thus, these results further highlight the functional diversity within these subtypes: Vglut2+ axons displayed a greater response to aversive stimuli than rewards, Calb1+ axons displayed greater response to rewards than aversive stimuli and was robustly sensitive to reward size, while reward responses were not detectable from Anxa1+ axons, which instead displayed a signaling decrease to aversive stimuli.

### Dopamine neuron functions differentially map onto genetic subtypes

To explicitly demonstrate the connection between functional and genetic dopamine neuron subtypes, we plotted the locomotion signaling (PC1/PC2 angles from Fig. 2L), response to rewards and response to air puffs for the subset of recordings where all 3 measurements were made (Fig. 5A). Calb1+, Vglut2+ and Anxa1+ recordings resided in separable regions of this 3D functional space, with minimal overlap. We then asked whether an unsupervised classification method, k-means clustering, could distinguish the subtypes based on these functional dimensions. Indeed, when searching for 3 clusters within the functional space, k-means separated the recordings into clusters that matched the genetic subtypes with 96% accuracy (Fig. 5B; of note, random chance = 33% accuracy; 100% accuracy for Vglut2+, 92% for Calb1+, and 96% for Anxa1+). Thus, our findings establish a clear connection between functional responses and genetic dopamine neuron subtypes and demonstrate that genetically defined subtypes of striatonigral dopamine axons have, on average within a small recording volume, markedly different signaling patterns during locomotion, reward and aversive stimuli.

**Figure 5.**
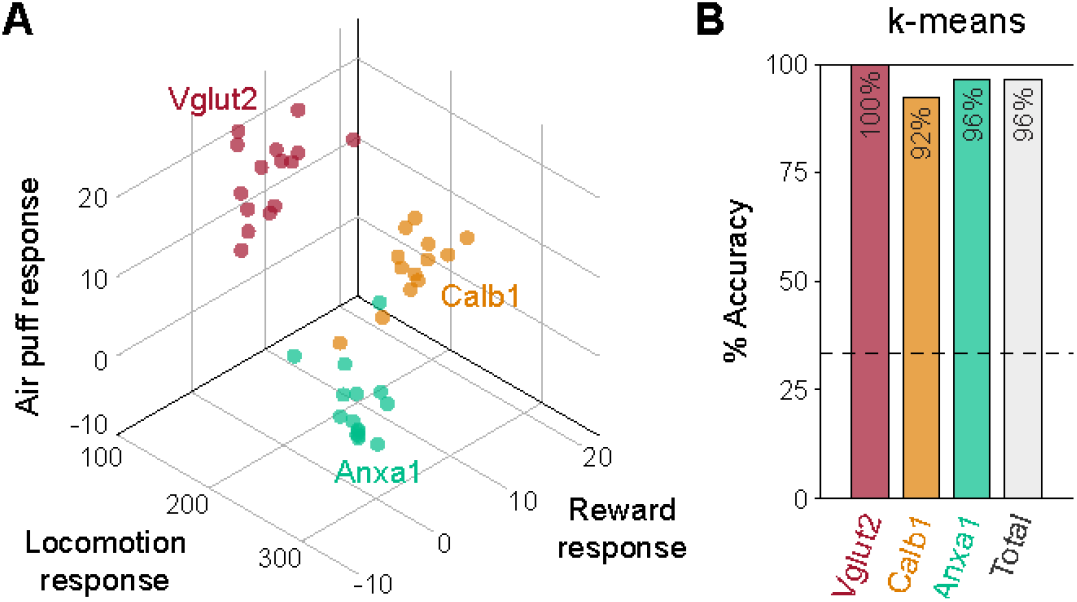
Unique locomotion, reward and air puff responses differentially map onto genetic subtypes of dopamine neurons. (A) 3D plot showing locomotion (PC1/PC2 angle, see Fig. 2L), reward and air puff responses for each recording and each subtype. (B) Unsupervised k-means classification distinguished subtypes based on locomotion (PC1 and PC2 scores), reward, and air puff responses, with total accuracy of 96%: 15/15 Vglut2, 12/13 Calb1 and 27/28 Anxa1 recordings correctly classified. Dashed line represents chance accuracy (33%).

### Axons track somatic signaling within subtypes

Slice studies have shown that coordinated activation of striatal cholinergic interneurons can not only modulate, but also trigger dopamine release in the absence of somatic firing^41,42,50,51^. A pioneering *in vivo* study has recently provided strong support for the idea that this local mechanism plays a significant role in dopamine release during behavior^52^. This study found that dopamine release from striatal axons co-varied with reward expectation, while firing in the midbrain somas did not, and further observed fast striatal dopamine release during certain behavioral epochs that did not correspond with somatic firing. However, establishing that dopamine is released from axons independently of somatic firing *in vivo* requires that axonal and somatic recordings are made from the same neurons^53^. Thus, an alternative explanation for any observed soma-axon signaling differences is that the striatal dopamine detected was released by a different set of axons than those belonging to the recorded somas–an experimental recording problem that could be rectified by labeling and recording from only one genetic subtype at a time.

Given the functional differences we observed in axons of different subtypes, we first asked whether the somas of these same subtypes show similar functional signaling as their axons. We repeated the above photometry recording experiments, but placed the optic fiber in SNc instead of striatum. At somas, GCaMP transients are caused by somatic action potential firing. Indeed, just as in the axonal recordings, Calb1+ and Vglut2+ somas responded to rewards and air puffs, while no detectable reward response was found in Anxa1+ somas (Fig. S7A-C). Calb1+ somas also showed greater responses to rewards and air puffs, and Vglut2+ somas on average had greater responses to air puffs than rewards (Fig. S7D), as their axons did. Calb1+ somas also showed greater responses to larger rewards (Fig. S7E). Furthermore, soma recordings from each of the subtypes showed highly similar signaling during locomotion compared to axons (Fig. S7G-I), and fell into the same, separable regions of the 3D functional space as axonal recordings of the same subtypes (Fig. S7J). Thus, axons and somas of the same dopamine neuron subtype displayed highly similar signaling correlations to locomotion and responses to rewards and aversive stimuli. This is further evidence that functional responses map onto genetic subtypes, as somas of individual subtypes intermingled to a fair degree in SNc, particularly within the photometry recording volume.

However, it is still possible that somas and axons could have similar correlation to movements or stimuli, but low correlations to each other (for example, somas and axons could be active at different accelerations or stimuli). Therefore, we performed simultaneous striatal axon and SNc soma recordings. Before recording from dopamine neuron subtypes, we first asked whether we could reproduce the soma-axon signaling differences previously described in non-subtype specific recordings ^52^, but with GCaMP and in head-fixed mice running on a treadmill. We labelled non-subtype-specific dopamine neurons (DAT mice) and used fiber photometry to simultaneously record from populations of axons in the striatum with one fiber and SNc somas with another fiber (Fig. 6A-B). We recorded from a range of random locations within striatum and SNc and often observed highly dissimilar signaling (ΔF/F) between striatal axons and SNc somas (Fig. 6C). Accordingly, the mean cross-correlation between axonal and somatic ΔF/F traces (Fig. 6D-E) was 0.37, which is a relatively low correlation for traces that have similar temporal dynamics (in contrast to cross correlations between ΔF/F and accelerations where the traces have dissimilar temporal dynamics, Fig. 2F). Therefore, similarly to previous reports^52^, we found somatic and axonal dopamine neuron signaling that was often very different when dopamine neurons are indiscriminately labelled.

**Figure 6.**
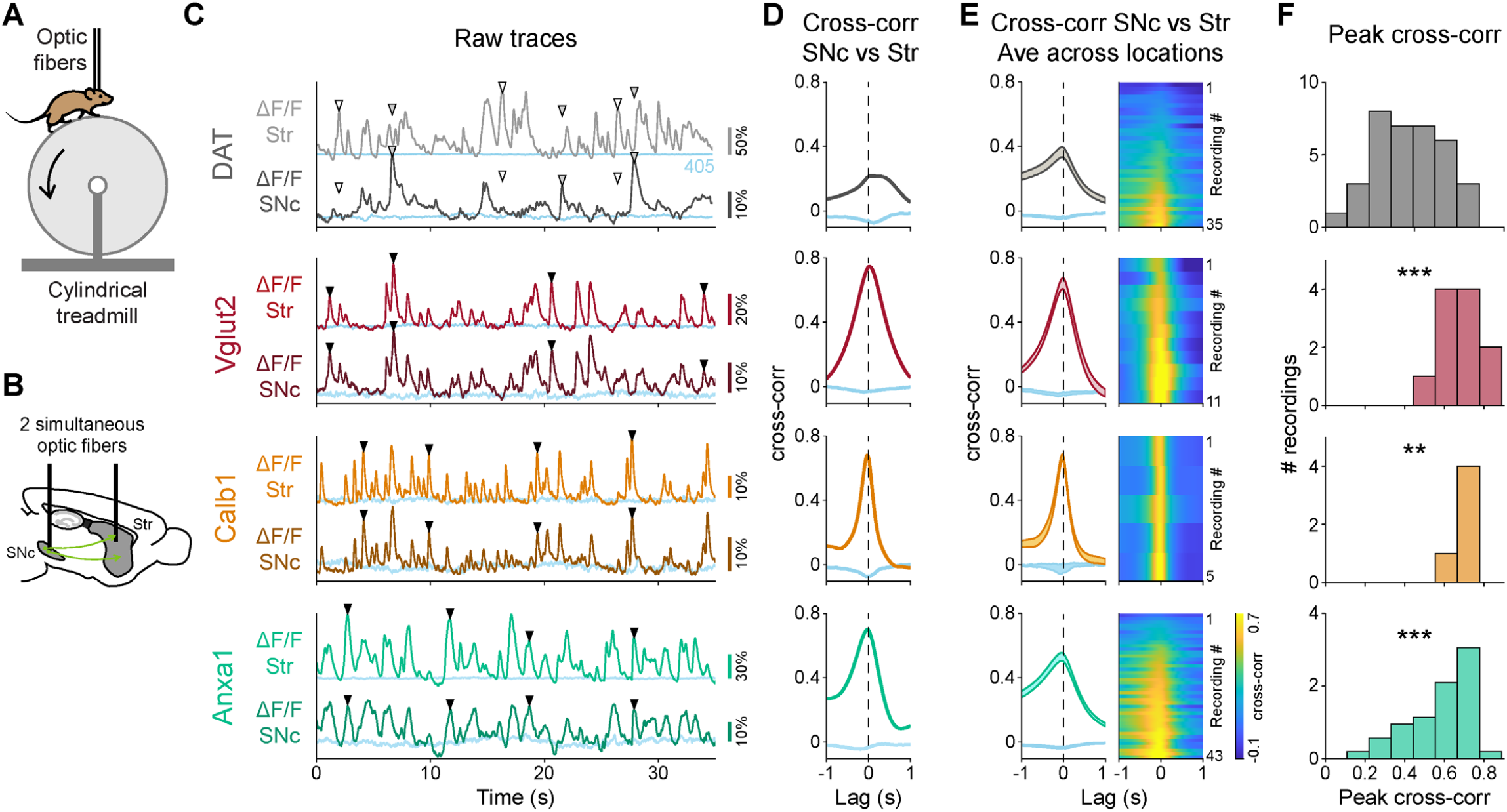
Highly-correlated signaling in axons and somas within genetic subtypes of dopamine neurons. (A) Mouse running on treadmill during dual fiber photometry. (B) Schematic of simultaneous photometry recordings from SNc and striatum. (C) Example recordings for DAT and each subtype showing simultaneous fluorescence traces (ΔF/F) from SNc and striatum. Isosbestic controls in blue. ▾= Example transients present in SNc and in striatum, ▽= example transient present in striatum but not in SNc (white fill) or vice-versa (gray fill). (D) Cross-correlation between ΔF/F traces from striatum and SNc shown in C. Isosbestic controls in blue. (E) Average cross-correlation between simultaneous ΔF/F traces from striatum and SNc for all recordings of each subtype and DAT. Isosbestic controls in blue. Shaded regions denote mean ± s.e.m. Heatmap shows cross correlations for each paired recording sorted by peak magnitude. DAT mice = 5, n = 35 recordings; Vglut2 mice = 4, n = 11; Calb1 mice = 2, n = 5; Anxa1 mice = 8, n=43. (F) Distribution of peak cross correlations between SNc and striatum for recordings of all subtypes and DAT shown in E. P-values for comparison to DAT: Vglut2 = 3×10-04, Calb1 = 3×10-03, Anxa1 = 3 x10-04 (Mann-Whitney U test with Bonferroni correction).

In contrast, when we repeated these soma-axon recordings from isolated subtypes, we found highly similar signaling between striatal axons and SNc somas (Fig. 6C), resulting in high cross-correlations (Fig. 6D), and this was consistent across recordings (Fig. 6E). On average, the cross-correlation between soma and axon ΔF/F recordings was significantly higher compared to DAT+ recordings (Fig. 6E-F; mean = 0.65 for Vglut2+, 0.67 for Calb1+, 0.58 for Anxa1+, compared to 0.37 for DAT; p-values for comparison with DAT+ = 3×10^−04^ for Vglut2+, 0.003 for Calb1+, 3×10^−04^ for Anxa1+, Mann-Whitney U test with Bonferroni correction). Overall, we conclude that recording from isolated dopaminergic functional subtypes leads to highly similar signaling patterns between somas and axons in behaving mice.

## Discussion

Here, we first found functional heterogeneity within the well-known Aldh1a1+ subtype, which motivated our use of single-nucleus transcriptomics to refine the existing classification of dopamine subtypes and led to the discovery of a new subtype characterized by Anxa1 expression within the previously described SNc Aldh1a1+ subtype. We then isolated and recorded from this new Anxa1+ subtype, as well as the known Calb1+ and VGlut2+ subtypes, and found unique functional signaling patterns to rewards, aversive stimuli, accelerations and decelerations. We made three main findings. i. While the Calb1+ and VGlut2+ subtypes robustly respond to unexpected rewards and aversive stimuli, such responses were not detected in the Anxa1+ subtype, even at striatal locations where its axons overlapped with the other subtypes. ii. Acceleration-and deceleration-correlated responses were differentially observed in genetically distinct neurons. The unique functional signaling differences between subtypes were also observed in recordings from SNc somas. iii. When dopaminergic subtypes were genetically separated, soma signaling correlated well with axonal signaling. These findings establish a connection between functional responses and genetic subtypes of dopamine neurons across a range of functional dimensions, validating the behavioral relevance of molecular classification schemes.

Though we here found significant differences in functional responses between SNc dopamine subtypes across different midbrain and striatal regions, fiber photometry records the mean fluorescence signal from populations of axons or cell bodies in the ∼300 micron recording volume. Thus, it is possible that some heterogeneity exists within the genetic subtypes at the single-cell/axon level. Nonetheless, similar signaling patterns to our reported averages have been observed in single-cell^11–13,17–21^ and single-axon^16^ recordings. This suggests that the functional differences we observed between subtypes are due to the strong enrichment of particular functions at the single cell level for specific subtypes. Thus, genetic subtypes provide a tool to reproducibly access different dopamine neuron functions, which is particularly important given the literature’s many conflicting observations/hypotheses on the role of dopaminergic neurons.

While the general assumption has been that all midbrain dopamine neurons respond to unexpected rewards, there has been scattered evidence against this dogma. A few single cell studies reported some SNc dopamine neurons that did not respond to rewards^17,54^, and axonal imaging recordings in dorsal striatum found several single axons not encoding rewards^16^. However, other studies have found reward responses in similar regions^15,23,24^. Since we detected robust reward responses in Calb1+ and Vglut2+ neurons, but not in Anxa1+ neurons, and since these different subtypes have different midbrain distributions and striatal projection targets, our results may help explain the previous discrepancies; different subtype(s) may have been investigated based on the recording location in SNc or striatum. Further, our functional characterization of Vglut2+ neurons agrees with previous recordings from overlapping soma/axon regions that reported aversive stimuli and reward signaling ^11–13^, with insensitivity to reward-size^12,55^. Based on these properties, such neurons have been proposed to signal novelty or salience^11,13^, or to reinforce avoidance of threatening stimuli^12^. Thus, of the three subtypes studied here that account for most SNc dopamine neurons, only Calb1+ neurons displayed robust reward size sensitivity, a hallmark of reward prediction error and involvement in positive reinforcement learning ^48,49^.

Previous research has reported that many SNc dopamine neurons signal at accelerations during a variety of motor tasks, but with differences in whether the neurons increase or decrease their firing at accelerations^16–21,56^. Since here we found that such signaling patterns were differentially expressed by the different subtypes, and since their cell body and axon locations are anatomically biased, these previous discrepancies might also be explained by the unknowing recording of different subtype(s) across studies based on location. For example, recordings in more medial SNc/lateral VTA (Calb1+ location) found most neurons decrease their firing at accelerations and respond to rewards^19^; recordings from dorsal striatum axons (Anxa1+ axon location) found increases in signaling at accelerations and no detectable reward responses^16^; and recordings from a broader range of locations (and thus subtypes) in SNc found neurons with both increases and decreases of firing at accelerations^17^–all of which agree with our results when considering subtype anatomical distributions.

Numerous hypotheses have been proposed to explain the function of fast dopamine signaling during locomotion: some suggest they increase the probability of movement initiations or the vigor of movements^17,57^, while others propose they function as a corollary discharge signal associated with particular reward-tied actions and involved in credit assignment^6^ or motor learning^58,59^. Again however, these differences in results and interpretations may lie in which dopamine neuron subtypes were recorded or manipulated in previous studies–for example, the former idea is supported by the optogenetic activation of dorsal striatum axons^16^ (likely Anxa1+ axons), while the latter is supported by studies of medial SNc and lateral VTA neurons^6^ (likely Calb1+ somas). Future optogenetic perturbation studies focused on the specific subtypes described here should help to provide further understanding of their role in behavior. Such research will also need to consider that many dopaminergic neurons co-release other neurotransmitters–Vglut2+ neurons co-release glutamate^60^ and Aldh1a1+ neurons may co-release GABA^61,62^ (though see^61^)–which likely play additional functional roles within striatum^60,63^. Importantly, however, previous research using fluorescent dopamine sensors indicate that, at least when such recordings are targeted to regions dominated by a particular subtype such as the dorsal striatum^64^, dopamine release is consistent with that expected based on the GCaMP transients reported here.

When the diversity of dopaminergic neurons was taken into account, we found high correlation between somatic and axonal signaling. This is consistent with the classical view that striatal dopamine release is controlled by anterogradely propagating action potentials originating in midbrain somas, rather than by local striatal modulation controlling dopamine release. This finding is also in agreement with previous reports demonstrating that cholinergic interneurons and dopamine axons in striatum are often desynchronized during behavior^65^, making it difficult to explain the majority of dopamine release based on local cholinergic control. However, this does not exclude the possibility that local cholinergic modulation may still play a role in controlling dopamine release at specific behavioral time points. For example, striatal dopamine and acetylcholine signaling have been found to synchronize at certain times during behavior, such as at locomotion initiation or during turning^42,65^. Regardless, our results here provide evidence that axons track somatic signaling within dopaminergic subtypes, indicating that such subtypes should be considered in order to fully understand the mechanisms of dopamine release in striatum during behavior.

Finally, our results provide new potential research directions for different dopamine related diseases, such as Parkinson’s disease, since there is emerging evidence that several of the subtypes studied here exist in humans^44^. The cell body locations and axonal projections of Aldh1a1+ match the pattern of dopamine loss in Parkinson’s disease^66–69^, and these neurons are especially vulnerable in Parkinson’s disease^43,44^, for which the Aldh1a1+ subtype has garnered considerable attention^44,58,70–73^. This highlights the importance of the functional heterogeneity found here, which led to our discovery of the new Anxa1+ subtype. Given the functional properties of these Anxa1+ neurons (acceleration-correlated signaling but no detectable reward response), this subtype may be particularly important in the context of Parkinson’s disease.

## Acknowledgments

We thank Hailey Kim for her help with data collection

## Author contributions

Conceptualization: MA, ZG, RA, DAD

Formal analysis: MA, ZG, RA, DAD

Investigation: MA, ZG, CD

Resources: CR, LF, YSK, KD

Visualization: MA, ZG

Supervision: RA, DAD

Writing: MA, ZG, RA, DAD

## Materials and Methods

### Animals

All animals used in this study were maintained and cared following protocols approved by the Northwestern Animal Care and Use Committee. Cre mouse lines were maintained heterozygous by breeding to wild-type C57BL6 mice. The Th-Flpo line and the Ai93D reporter line were maintained homozygous. The Dat-tTA mouse line was maintained heterozygous by breeding with the Ai93D reporter. The Aldh1a1-iCre and Anxa1-iCre lines was generated at Northwestern University by the Transgenic and Targeted Mutagenesis Laboratory.

Both males and females were used for all experiments. Adult mice were used for viral injections at 2 to 4 months old. For indiscriminate labelling of SNc dopamine neurons, DAT-IRES-Cre mice (RRID: IMSR_JAX:027178) were injected with AAV1-CAG-FLEX-GCaMP6f virus (RRID: Addgene_100835). For labelling of SNc Anxa1+ neurons, Anxa1-iCre mice (new line) were injected with AAV1-CAG-FLEX-GCaMP6f virus (RRID: Addgene_100835). For labelling VGlut2+ or Aldh1a1+ dopamine neurons, we crossed VGlut2-IRES-Cre (RRID: IMSR_JAX:016963) or Aldh1a1-iCre mice (new line) with Th-2A-Flpo mice^23^, and offspring were injected with AAV8-EF1α-CreOn/FlpOn-GCaMP6f virus (RRID: Addgene_137122). For labelling Calb1+ dopamine neurons, we crossed Calb1-IRES2-Cre mice (RRID: IMSR_JAX:028532) with DAT-tTA (RRID: IMSR_JAX:027178), Ai93D (CreOn/tTAOn GCAMP6f reporter) (RRID: IMSR_JAX:024107) mice.

### Generation and characterization of the Aldh1a1-iCre line

Because our previous Aldh1a1-CreERT2 strain displayed substantial mosaicism, resulting in only weak GcaMP6f signals, we opted to generate an Aldh1a1-iCre strain (Fig. S4A). The Aldh1a1-iCre line was generated at Northwestern University by the Transgenic and Targeted Mutagenesis Laboratory. In brief, a P2A peptide directly followed by iCre and a BGH polyA sequence were inserted after the last encoded amino acid of Aldh1a1, using CRISPR mediated HDR (Guides 1-2 ACGACTATGCTGGTTAC, TCCCCCTTTAGGGG TGAGCA). First, PRXB6/N ES cells were electroporated and screened for insertion and correct locus with multiple primer pairs (Aldh1a1-iCre insertion primers Forward 1-3 CTATTCACTTGCAGTTGGCTTGG, GCTTGGAGGTTTCCTAAGTGTG, AGATCCCTGATGGAGAACTCTG and Reverse 1-3 GTCCAGGGTTCTCCTCCACG, GCATGATTTCAGGGATGGACAC, CATCCTTGGCACCATAGATCAG) followed by Sanger sequencing of iCre+ clones from outside the homology arms through the construct in order to confirm fidelity of the insertion. Clone C7 was expanded and injected into blastocysts to generate chimeras and used for all experiments herein. Aldh1a1-iCre mice were genotyped using primer set 3 described above. To determine the expression fidelity of this allele, 0.4 µl of AAV5-EF1α-DIO-mCherry (RRID:Addgene_37083) was injected into SNc bilaterally (coordinates relative to bregma: x = ±1.45mm, y = -3.15mm, z = -3.1, -4.1, -4.4, -4.7mm, 0.1 µl at each depth) in n = 4 adult mice. Three weeks later, mice were perfused, and brains were sectioned at 25 µm for immunofluorescence staining. Floating sections were first blocked for 24 hours at 4°C in PBS containing 0.03% Triton-X and 5% normal donkey serum. Sections were incubated with primary antibodies against Aldh1a1 (goat, R&D Systems Cat# AF5869, RRID:AB_2044597), TH (mouse, Sigma-Aldrich Cat# T2928, RRID:AB_477569 ; Pel-Freez Biologicals Cat# P40101-0, RRID:AB_461064) and mCherry (rat, Thermo Fisher Scientific Cat# M11217, RRID:AB_2536611) in blocking buffer for 24 hours, followed by 4 washes in PBS-Tween20 and incubation with secondary antibodies (Donkey anti Goat Alexa Fluor 488 [Molecular Probes Cat# A-11055, RRID:AB_2534102], Donkey anti Mouse Alexa Fluor 647 [Thermo Fisher Scientific Cat# A-31571, RRID:AB_162542], Donkey anti Rabbit Alexa Fluor 647 [Thermo Fisher Scientific Cat# A-31573, RRID:AB_2536183], Donkey anti Rat Cy3 [Jackson ImmunoResearch Labs Cat# 712-165-153, RRID:AB_2340667], and DAPI [Thermo Scientific 62248]) for 2 hours at room temperature. Sections were then imaged at 20x. For each brain, 4-5 sections spaced at least 100 microns apart and centered about the area of maximal viral recombination were counted for mCherry+/DAPI+/Aldh1a1+ and mCherry+/DAPI+/Aldh1a1-cells (2740 cells total) (Fig. S4B, C).

### Generation and characterization of the Anxa1-iCre line

To access the Anxa1+ dopamine neurons, the Anxa1-iCre line (Fig. S4D) was also generated by the Transgenic and Targeted Mutagenesis Laboratory, using similar methodologies as above. For CRISPR mediated HDR, Guides 3-4 (AAGATTCTGGTGGCCCTCTG, ACTTAAGCCCATGCCAT) were used. Clones were screened for insertion using iCre genotyping primers (primer set 3 from Aldh1a1-iCre insertion validation described above). To determine the expression fidelity of this allele, 0.4 µl of AAV5-EF1α-DIO-mCherry (RRID: Addgene_37083) was also injected into SNc bilaterally at the same coordinates as above, and stained for immunofluorescence using the same protocol as above, but with Rabbit anti Anxa1 antibody (Thermo Fisher Scientific Cat# 71-3400, RRID: AB_2533983) in place of Aldh1a1. Sections were then imaged at 20x. Viral recombination occurred in cells with both high Anxa1 expression and faint Anxa1 expression (Fig. S4E). To confirm that Anxa1-iCre recombination was limited to a subset within Aldh1a1-expression DA neurons, we also co-stained for Aldh1a1 and mCherry using the same antibodies as above, which showed many Aldh1a1+ cells that did not express the reporter (Fig. S4F). To corroborate this further, we stained Anxa1-iCre, TH-Flpo, RC::FrePe mice for Aldh1a1 and GFP (the expression of which is dependent on both iCre and Flpo recombination) using the same protocol as above, which showed Aldh1a1 expression to be broader than Anxa1-iCre expression in development, thus confirming that our viral labeling results were not an artifact of insufficient viral delivery and/or diffusion(Fig. S4G).

### Integration of single-cell RNAseq datasets

Data was analyzed using custom R code (in RStudio).

Data from four prior single-cell studies (Tiklova et al. 2019 (GEO: GSE116138), Saunders et al. 2018 (Dropviz.org), Kramer et al. 2018 (GEO: GSE115070), and La Manno et al. 2016 (https://bioconductor.org/packages/release/data/experiment/html/scRNAseq.html)) was acquired for integration using Seurat version 3.2.0.. For Saunders et al. data, specific clusters identified as TH+ substantia nigra neurons (SN clusters 4-1, 4-2, 4-3, 4-4, 4-5, 4-6, 4-7, 4-8, 4-9, & 3-7) were subsetted and used integration. Violin plots of number of reads and number of genes for each dataset were generated and used to determine cutoffs for pre-filtering of each dataset prior to integration to remove doublets or low-quality cells (Fig S2C).The following filters were ultimately applied: Saunders – nFeatures < 3500, mitochondrial read % <25, nCount < 10,000. Tiklova -- nFeatures > 6000, mitochondrial read % < 10, nCount < 3500000. Kramer 00 nFeatures > 5000, nCount > 400000. Datasets were normalized individually and integrated using the recently described SCTransform pipeline (Hafemeister & Satija 2019 PMID: 31870423) with default settings and regression on percent mitochondrial reads. Principle component analysis was performed on the subsequent integrated dataset, and an elbow plot was used to determine the number of PCs used for clusters (18 PCs were ultimately used). Clustering was performed using the standard Seurat pipeline at default settings, resulting in 8 clusters (Fig S2A). Determination of marker genes for clusters was performed using the FindAllMarkers command in Seurat on the RNA assay with the following settings: min.diff.pct = 0.20, only.pos=TRUE, min.pct = 0.05). Of note, exploring differential expression of marker genes using a heatmap (not shown) revealed a unique signature distributed across multiple clusters that did not appear to fit any other subtypes. Examining the source of these cells revealed they came entirely from the Tiklova et al. dataset. Therefore, to explore the potential inclusion of a unique group of cells stemming from that dataset, we re-clustered our dataset using the LIGER R package version 2.0.1, which differs from Seurat dataset integration in that it is designed to account for the potential inclusion of unique cell types stemming from only individual samples being integrated (Welch et al. 2019 PMID: 31178122). Clustering with LIGER revealed a cluster of distantly related cells which came entirely from Tiklova et al (Fig S2D). Due to the distinct signature of these cells which did not match the clusters they were placed in using the Seurat integration, these cells were subsequently filtered out of our dataset. After this, all clusters were represented by all source datasets (Fig S2E). Violin plots of the top 2 defining markers per cluster were generated using the Seurat VlnPlot command with default settings (Fig S2F).

### Single nucleus RNA sequencing

To isolate nuclei for snRNA-seq library generation, n=5 mice (3 female, 2 male) were sacrificed and rapidly decapitated for extraction of brain tissue. A 2-3mm thick block of ventral midbrain tissue was dissected out and collected for subsequent isolation. Tissue was dounce homogenized in a nuclear extraction buffer (10mM Tris, 146mM NaCl, 1mM CaCl2, 21mM MgCl2, 0.1% NP-40, 40u/mL Protector RNAse inhibitor (Roche 3335399001). Dounce homogenizer was washed with 4mL of a washing buffer (10mM Tris, 146mM NaCl, 1mM CaCl2, 21mM MgCl2, 0.01% BSA, 40U/mL Protector RNAse inhibitor) and filtered through a 30uM cell strainer. After three rounds of washing by centrifugation (500g for 5 minutes) and resuspension in a nuclei resuspension buffer (10mM Tris, 146mM NaCl, 1mM CaCl2, 21mM MgCl2, 2% BSA, 0.02% Tween-20), nuclei suspension was stained with DAPI and filtered through a 20uM strainer. This nuclei suspension was then sorted via FACS with a 100uM nozzle at a frequency of 30.0K and pressure of 20 PSI, with gates set for isolation of GFP+ singlet nuclei (Fig S3A). A total of 50,500 nuclei were sorted across all samples, which was subsequently used for preparation of two 10X Genomics Chromium libraries (one for pooled male mice, one for pooled female mice).

Library preparation was performed by the Northwestern University NUSeq Core Facility. Nuclei number and viability were first analyzed using Nexcelom Cellometer Auto2000 with AOPI fluorescent staining method. Sixteen thousand nuclei were loaded into the Chromium Controller (10X Genomics, PN-120223) on a Chromium Next GEM Chip G (10X Genomics, PN-1000120), and processed to generate single cell gel beads in the emulsion (GEM) according to the manufacturer’s protocol. The cDNA and library were generated using the Chromium Next GEM Single Cell 3’ Reagent Kits v3.1 (10X Genomics, PN-1000286) and Dual Index Kit TT Set A (10X Genomics, PN-1000215) according to the manufacturer’s manual with following modification: PCR cycle used for cDNA generation was 16 and the resulting PCR products was size-selected using 0.8X SPRI beads instead of 0.6X SPRI beads as stated in protocol. Quality control for constructed library was performed by Agilent Bioanalyzer High Sensitivity DNA kit (Agilent Technologies, 5067-4626) and Qubit DNA HS assay kit for qualitative and quantitative analysis, respectively.

The multiplexed libraries were pooled and sequenced on Illumina Novaseq6000 sequencer with paired-end 50 kits using the following read length: 28 bp Read1 for cell barcode and UMI and 91 bp Read2 for transcript. Raw sequence reads were then demultiplexed and transcript reads were aligned to mm10 genome using CellRanger with --include-introns function.

### Analysis of single nucleus RNA sequencing data

Data was analyzed using custom R code (in RStudio).

Outputs from CellRanger were read into Seurat version 4.0.2 using the Read10X command for each sample. Numbers of UMIs, features and mitochondrial reads were plotted for each dataset (Fig S3C) and used to determine cutoffs for quality control pre-filtering of each sample; nuclei with fewer than 500 unique features were removed from each dataset. The male and female datasets were then normalized and integrated using the SCTransform V2 pipeline (Choudhary & Satija, 2022 PMID: 35042561) using all default settings and regression on percent mitochondrial reads. In total, the integration resulted in a final dataset of 12,065 nuclei, with a mean UMI count of 3435 and mean of 1683 features. Clustering was performed using the Seurat FindClusters command using 30 principle components and a resolution of 0.5. Differential expression tested was performed using the FindAllMarkers command on the SCT assay with default settings, with the exception of logfc.threshold = 0.15 in order to better detect differential expression of genes with low overall detection rates in the dataset. Determination of Sox6+ and Calb1+ significant clusters was made using a Wilcoxon Rank Sum Test by running the FindAllMarkers Seurat command with the following settings: features = c(“ Sox6”, “ Calb1”), min.pct = 0, min.diff.pct = 0, logfc.threshold = 0, only.pos = TRUE.

In order to better visualize the expression of marker genes, we then performed zero-preserving zero-imputation using ALRA (Linderman et al., 2022 PMID: 35017482), which aims to increase the detection of low-expression genes while preserving true biological zeros. Zero-imputed data was used solely for visualizations of features as seen in Figures 1B, 1C, S3F and S3I, but not for any statistical determination of differential expression. Heatmap of the top 4 marker genes for each cluster (Fig S3H) was generated using the top 4 differentially expressed genes (determined per average log fold change) for each cluster, filtering for only unique genes.

### Stereotaxic viral injections

Adult mice (postnatal 2-4 months old) were anesthetized with isoflurane (1-2%), and a 0.5–1-mm diameter craniotomy was made over the right substantia nigra (−3.25 mm caudal, +1.55 lateral from bregma). A small volume (0.4 μl total) of virus (AAV8-EF1α-CreOn/FlpOn-GCaMP6f (RRID:Addgene_137122, titer 6.10E+13) for Aldh1a1-iCre/Th-Flpo and VGlut2-IRES-Cre/Th-Flpo mice, or AAV1-CAG-FLEX-GCaMP6f (RRID:Addgene_100835, titer 2.00E+13) for DAT-Cre or Anxa1-iCre mice), diluted 1:1 in PBS, was pressure injected through a pulled glass micropipette into the SNc at 4 depths (−3.8, -4.1, -4.4 and -4.7 mm ventral from dura surface, 0.1 μl per depth). Following the injections, the skull and craniotomy were sealed with Metabond (Parkell) and a custom metal headplate was installed for head fixation. The location of recording sites was marked the surface of the Metabond for future access. For Calb1-IRES2-Cre/DAT-tTA/Ai93D mice, which express GCaMP6f endogenously, no injection was conducted and only the headplate was implanted at this time. 4 weeks were allowed for GCaMP6f expression to ramp up and fill dopaminergic somas in SNc and axons in striatum.

### Training and behavior

Starting 1-2 weeks after injection, mice were head-fixed with their limbs resting on a 1D cylindrical Styrofoam treadmill ∼20 cm in diameter by 13 cm wide in the dark. Mice were habituated on the treadmill for 3-10 days until they ran freely and spontaneously transitioned between resting and running. Rotational velocity of the treadmill during locomotion was sampled at 1,000 Hz by a rotary encoder (E2-5000, US Digital) attached to the axel of the treadmill and a custom LabView program.

After mice ran freely, a subset of mice were water restricted and received unexpected water rewards, aversive air puffs, and light stimuli while on the treadmill, using a custom LabView program. Large (16 μl) and small volume (4 μl) water rewards were delivered through a waterspout gated electronically through a solenoid valve, which was accompanied by a short ‘click’ noise. Air puffs were delivered by a small spout pointed at their left whiskers, which was connected to a ∼20 psi compressed air source and triggered electronically through the opening of a solenoid valve for 0.2s. Triggering of this solenoid was also accompanied by a ‘click’ noise. For light stimuli, a blue LED placed ∼30 cm in front of the head-fixed mouse was electronically triggered for 0.2s. Rewards, air puffs and light stimuli were alternated at random during recordings and delivered at pseudo-random time intervals (10-30s between any two stimuli).

### Fiber photometry

4 weeks after injection, mice were once again anesthetized, and a small craniotomy (1 mm in diameter) was drilled through the Metabond and skull, leaving the dura and cortex intact. Craniotomies were made at different locations depending on the experiment, which were pre-marked during the injection surgery: for SNc -3.25 mm caudal, +1.55 lateral from bregma, and different locations over striatum (for example -1.1 mm caudal +2.8 lateral, or +0.5 mm caudal + 1.8 lateral). The craniotomies were then sealed with Kwik-Sil (World Precision Instruments KWIK-SIL).

After the mice recovered from this short (10-15 min) surgery for one day, they were head-fixed on the linear treadmill, and the Kwik-Sil covering the craniotomies was removed. One or two optical fibers (200 μm diameter, 0.57 NA, Doric MFP_200/230/900-0.57_1.5m_FC-FLT_LAF) were lowered slowly (5 μm/s) using a micromanipulator (Sutter MP285) into the brain to various depths measured from the dura surface. In the striatum, recording depths ranged from 1.6 to 4.1 mm; in SNc, depths ranged from 3.5 to 4.5 mm. Recordings started at 1.6 mm in striatum, and 3.5 mm in SNc, but if no ΔF/F transients were detected at those depths the fiber was moved down in increments of 0.25-0.5 mm in striatum or 0.15-0.2 mm in SNc, until transients were detected. From there, a 15 min recording was obtained, and the fiber was moved further down in the same increments. Subsequent recordings were obtained until a depth was reached where transients were no longer detected, at which point the fiber was pulled out of the brain slowly (5 μm/s).

A custom-made photometry setup was used for recording. Blue excitation (470 nm LED, Thor Labs M70F3) and purple excitation light (for the isosbestic control) (405 nm LED, Thor Labs M405FP1) were coupled into the optic fiber such that a power of 0.75 mW emanated from the fiber tip. 470 and 405 nm excitation was alternated at 100 Hz using a waveform generator, each filtered with a corresponding filter (Semrock FF01-406/15-25 and Semrock FF02-472/30-25) and combined with a dichroic mirror (Chroma Tech Corp T425lpxr). Green fluorescence was separated from the excitation light by a dichroic mirror (Chroma Tech Corp T505lpxr) and further filtered (Semrock FF01-540/50-25) before collection using a GaAsP PMT (H10770PA-40, Hamamatsu; signal amplified using Stanford Research Systems SR570 preamplifier). A Picoscope data acquisition system was used to record and synchronize fluorescence and treadmill velocity at a sampling rate of 4 kHz.

### Histology

Immediately after the last recording, mice were perfused transcardially with PBS (Fisher) then 4% paraformaldehyde (EMS). Brains were stored in PFA at 4 °C overnight then transferred to 40% sucrose (Sigma) for at least 2 days before sectioning. Coronal slices (50 μm thick) were cut on a freezing microtome and stored at 4 °C in PBS. For immunostaining of dopaminergic neurons, sections were washed in PBS, blocked in PBS + 0.3% Triton-X (Sigma) + 5% normal donkey serum (Sigma), incubated overnight with primary antibodies Sheep anti-Tyrosine Hydroxylase (1:1000 dilution, RRID:AB_461070) and Rabbit anti-GFP, which recognizes GCaMP6f (1:1000 dilution, RRID:AB_221569), washed again in PBS + 0.3% Triton-X, then incubated with secondary antibodies tagging Tyrosine Hydroxylase with Alexa Fluor 555 (Donkey anti-Sheep Alexa Fluor 555, RRID:AB_2535857) and GCaMP6f with Alexa Fluor 488 (Donkey anti-Rabbit Alexa Fluor 488, RRID:AB_2313584). Images of SNc and Str were acquired on an Olympus or Keyence Slide Scanner (VS120 or BZ-X810, respectively) for verification of injection accuracy and fiber placement. Other brains were mounted and imaged without immunostaining for fiber placement. Histology is not available for 2 out of 14 DAT mice and 1 out of 5 Calb1 mice, and fiber tracks could not be identified for 2 out of 16 VGlut2 mice.

### Photometry data preprocessing

Analysis was conducted using custom code on MATLAB.

Simultaneous traces (velocity from rotary encoder, trigger signals for reward, air puff, and light stimuli delivery, licking from a lick sensor, fluorescence detected by PMTs from one or two optic fibers, and output from waveform generator used to alternate 405 and 470 nm illumination every 10 ms) were collected at 4 kHz by a Picoscope6 data acquisition system. Fluorescence collected during 405 or 470 nm illumination (20 time bins for each pulse of 405 or 470 nm excitation) was separated using the binary output from the waveform generator. For each transition period between illumination sources, 5 time bins were excluded to remove transition times. Traces were then re-binned to 100 Hz by averaging every 40 time bins for velocity and every 40 time bins for 405 and 470 fluorescents (but only including 15 of 40 bins for each source: excluding 20 bins when the alternate source was on and 5 transition bins).

We first corrected fluorescence traces for background signal (intrinsic fluorescence and any illumination bleed-through) by subtracting 85% of the baseline (baseline defined as 8^th^ percentile over a 20s window). This 85% was estimated from photometry recordings from cortex, which was unlabeled (no GCaMP expression), obtained from 10 recordings from 5 mice. 405 and 470 fluorescence traces were corrected independently. To calculate ΔF/F, traces were then normalized by baseline fluorescence division (8^th^ percentile over a 20s window) separately for 405 and 470. The subtraction and normalization steps together corrected for bleaching and removed any slow drifts in baseline. Next, traces were converted to ΔF/F units (baseline at 0) by subtracting the baseline (median of all non-transient bins for 470 nm traces, and median of all bins for 405 nm traces)

For comparison of traces between dopaminergic subtypes, ΔF/F traces were normalized so that the baseline remained at 0 and the largest transient peak for each trace was 100%. Throughout all figures, Norm ΔF/F units refer to this normalization (0-100 scale). 405 traces were normalized using the amplitude of the largest peak from the corresponding 470 traces. Example raw traces in Fig. 2C, 4C, 6C and Fig. S1D, S8A show non-normalized traces.

### Criteria for recording inclusion

Only recordings with signal-to-noise ratios greater than 10 were included in the analysis. To calculate signal-to-noise ratios for each recording, we selected well-isolated transients, as defined by having a large, fast rise (30 ΔF/F/s) immediately followed by a decay. We first removed all slow fluctuations except transients in (non-normalized) ΔF/F traces by subtracting the 8^th^ percentile over a window 2-3 times the width of observed ΔF/F transients (250 bins, 2.5s), and then smoothed the resulting trace over a 0.2s window (20 bins) to reduce noise. Transient rises and decays were identified by locating the zero-crossings on the derivative of the trace, also smoothed over 0.2s window. Only clearly isolated transients were included – those with a rise greater than 30 ΔF/F/s followed by a decay greater than -5 ΔF/F/s. Traces with less than 0.2 transients per second were excluded. Signal values for each recording were calculated as the 80^th^ percentile of isolated transient peaks. Noise for each recording was calculated by smoothing each (non-normalized) ΔF/F trace over 10 bins (0.1s), then subtracting this smoothed trace from the original ΔF/F trace and using the standard deviation of the resulting trace as the noise value. The signal and noise values were divided to obtain signal-to-noise for each trace. These steps for determining signaling to noise for each trace were not used for any further analysis.

ΔF/F traces from 405 nm illumination (isosbestic control) were used to remove any movement artifacts. While GCaMP6f fluorescence intensity is dependent on calcium concentration when excited with 470 nm light, it is still fluorescent but in a calcium-independent way when excited with 405 nm light (Tian et al. 2009). Therefore, calcium transients in neurons are detected with 470 nm illumination but are absent with 405 nm illumination, while movement artifacts are present in both traces. Movement artifacts were identified using the 405 nm traces from each recording as follows. (Non-normalized) 405 ΔF/F traces were smoothed over a 10-bin window (0.1s). This smoothed trace was subtracted from the original 405 ΔF/F trace, so that only the noise remained (same process as used above for 470 traces to separate noise and signal). A max noise value was calculated as the max absolute value of this noise trace. Any bins in the original 405 ΔF/F trace more than 3 times this max noise (or 3 times below -max noise) were excluded from further analysis. Additionally, any sequential bins that were above max noise (or below -max noise) for longer than 0.2 s (20 bins, less than half the width of observed calcium transients) were also excluded, with an additional 0.1s (10 bins) on both sides also excluded. Any bins removed from the 405 ΔF/F trace were also removed in the corresponding 470 ΔF/F and velocity traces. If more than 5% of the bins in a recording met these movement artifact exclusion criteria, the entire recording was excluded.

### Analysis of signaling during locomotion

Only locomotion time bins were included for locomotion analysis in Fig. 2, S1, S5, S7. Locomotion vs rest bins were selected using a double threshold on the velocity trace in both positive and negative directions (thresh1 = ± 0.024 m/s, thresh2 = ± 0.010 m/s). Isolated 1 bin-long locomotion periods (no other movement within 2 bins on either side) were excluded, as well as rest periods shorter than 0.5 s. Time bins were considered as locomotion periods only if they lasted longer than 0.5 s and had an average velocity greater than 0.2 m/s. For a recording to be included in the locomotion analysis, the recording needed to include a total of at least 100 sec of locomotion.

Acceleration was calculated from the velocity traces as the difference between consecutive treadmill velocity time bins (first smoothed over 6 bins, 0.06s), then multiplied by the sampling frequency (100 Hz) for proper m/s^2^ units.

Cross-correlations between ΔF/F and acceleration (Fig. 2D,F; 3A; S1E-F; S5D; S7G) were calculated for locomotion periods only (defined above) using MATLAB’s *crosscorr* function over a 1s lag window (100 time bins). The same process was used to calculate the cross-correlation between corresponding 405 ΔF/F traces and acceleration, and any recording with a peak cross-correlation (between 405 ΔF/F trace and acceleration) above 0.1 was excluded from all locomotion analysis.

For triggered averages of ΔF/F on accelerations and decelerations (Fig. 2G, S5E, S7H), isolated large accelerations and decelerations were selected by first locating the zero-crossings on the acceleration trace, considering individual accelerations/decelerations the interval between two zero-crossings of the trace. Accelerations/decelerations were included if they had a duration of at least 50 ms (0.05s) and a peak greater than 2 m/s^2^ (accelerations) or lower than -2 m/s^2^ (decelerations), but only if they were not surrounded by other large accelerations or decelerations (no acceleration > 2 m/s^2^ or < -2ms^2^ in a window of 0.25s on either side).

Conversely, for triggered averages of acceleration on ΔF/F transient peaks (Fig. 2H, S5F, S7I), we selected well-isolated transients from non-normalized ΔF/F traces, as defined by having a large, fast rise (30 ΔF/F/s) immediately followed by a decay (as used in the calculation of signal-to-noise ratio above).

For plotting of cross-correlation and triggered averages above, traces were smoothed over 5 time-lag bins (0.05s). Shaded areas represent the mean ± standard error of the mean (s.e.m.), while accompanying heatmaps show cross-correlations/triggered averages for all individual recordings. Heatmaps in Fig. S1F were sorted by the integral of the ΔF/F-acc cross-correlation at positive lags (see below), while heatmaps in Fig. 2F-H and Fig. S5D-F, S7G-I were sorted by PC1/PC2 angle (see PCA section below).

SNc recordings (Fig. S7G-J) were analyzed in the same manner as striatal recordings.

For analysis of timing differences between Calb1+ and VGlut2+ deceleration signaling shown in Fig. 2I, Fig. S5B-C, the lag between ΔF/F transient peaks and deceleration peaks was quantified by locating in time either the minimum cross-corr value between 0-1s for the ΔF/F-acceleration cross-correlations for each recording (Fig. 2I), the maximum ΔF/F value between 0-1s for the triggered average on deceleration (Fig. S5B), or the minimum acceleration value between -1-0s for the triggered average on transient peaks (Fig. S5C).

For the initial functional characterization shown in Fig. S1H-I, differences in locomotion signaling were quantified by calculating the integral of the cross-correlation between ΔF/F and acceleration at positive lags (0-1s), where positive values indicate a peak in the cross-correlation and thus ΔF/F transients following accelerations, while negative values indicate a trough and thus ΔF/F transients following decelerations. For the quantification of acceleration/deceleration signaling across depths in striatum shown in Fig. S1H (but also Fig. S5I), depth from surface as shown in was defined as the depth at which the fiber tip was located from the brain surface, as measured by the micromanipulator used to move the fiber during photometry. To reduce overlap between data points at the same depth plotted, a random amount between +0.1 and -0.1 mm was added to each depth. This measure of locomotion signaling was also used to plot the relationship between locomotion signaling and reward responses in Fig. S1I (for reward response calculation, see “ Analysis of responses to rewards and air puffs” section below), and to sort the ΔF/F-acceleration correlation plots in Fig. S1F.

### Principal component analysis (PCA) of locomotion signaling

Principal component analysis (PCA) was applied to the matrix of all cross-correlation traces from striatal recordings (shown in Fig. 2F), from all functionally homogeneous subtypes (VGlut2+, Calb1+ and Anxa1+), using MATLAB’s *pca* function (without centering: *‘Centered’, ‘off’*; however, equivalent results were obtained when we repeated the PCA analysis with centering, data not shown). This function outputs the principal components (loadings, eigenvectors), the scores for each recording’s cross-correlation along each principal component (matrix of all SNc cross-correlation traces multiplied by the loadings matrix), and the variance explained by each principal component across all recordings. For the representation of combinations of the first two principal components (PC1 and PC2) shown in Fig. 2J, PC1 and PC2 were weighted by the standard deviation of their scores across recordings (∼1 for PC1, ∼0.7 for PC2), to accurately represent each quadrant in Fig. 2K-L, S5G-H. Fig. 2K and Fig. S5G-H show the PC1 and PC2 scores for each recording of each subtype. In Fig. S5H, recordings were color-coded based on the depth from brain surface at which they were recorded, as measured by the micromanipulator used to move the fiber during photometry.

For SNc recordings, the cross-correlations between ΔF/F and acceleration for all recordings of all subtypes, as shown in Fig. S7G, were decomposed using the same principal components calculated above from the striatal cross correlations. Scores for SNc cross-correlations (Fig. S7J) were calculated by multiplying the matrix of all SNc cross-correlation traces by the striatal loadings matrix (principal components). The % of SNc variance explained by each principal component (PC1 = 53.2% of variance, PC2 = 24.3%) was calculated as the variance without the mean subtracted (not centered).

In the PC1/PC2 space shown in Fig. 2K, the angle of each point from the origin represents the shape of the cross-correlation between acceleration and ΔF/F, and thus the different relationships between subtypes’ signaling and acceleration, while the distance from the origin represents the amplitude of the cross-correlation. To quantify the shape of the cross-correlation across subtypes, we calculated the angle of each recording in the PC1/PC2 space (with each PC weighted by its standard deviation) and plotted it in a radial histogram (Fig. 2L). P-values for reporting statistical significance of the difference between subtypes across this PC1/PC2 space were calculating by opening the angular space at 45° (the region where the least recordings from Calb1/VGlut2/Anxa1 fall) and using a Wilcoxon rank-sum test with Bonferroni correction (multiply p-values by 3) to compare subtypes. This angle was also used to sort cross-correlation and triggered average heatmaps in Fig. 2F-H, S5D-F, S7G-I, starting by the middle of the quadrant opposite to the center of mass for each subtype. Fig. 3B show the location of each recording color-coded based on the PC1/PC2 angle for that recording. The colormap was defined by assigning a different color to each the middle of each quadrant (45°, 135°, 225°, 315°), where the center of mass of each subtype approximately falls at.

### Analysis of responses to rewards and air puffs

Reward delivery times were only included when the mice consumed the reward (detected by the lick sensor) within a 1s window from delivery. For analysis of rewards delivered at rest, rewards were excluded if there were any accelerations greater than 2.5 m/s^2^ (or decelerations greater than -2.5 m/s^2^) in a window of 0.75s before or after the reward delivery, or any accelerations greater than 1.5m/s^2^ (or decelerations greater than -1.5m/s^2^) within a 0.4s window after the reward (where responses to rewards are detected). Triggered averages on rewards (Fig. 4D-E, L, Fig. S1G, S6C-D, S7A), air puffs (Fig. 4F, L, Fig. S6EF, S7B), and rewards at rest (Fig. S6A) were calculated by averaging normalized ΔF/F traces (or licking traces for 4D, S6D) in a 1s window before and after included reward or air puff delivery times.

For plotting of triggered averages above, traces were smoothed over 5 time-lag bins (0.05s). Shaded areas represent the mean ± standard error of the mean (s.e.m.), while accompanying heatmaps show triggered averages for all individual recordings. Heatmaps in Fig. 4D-F and Fig. S1G; S6A, C-E; S7A-B were sorted by reward response size (see below).

To calculate the size of the response to each stimulus (change in fluorescence) shown in Fig. 4G-K; Fig. 5A and Fig. S1I, S6B, F, S7C-F, we calculated the difference between the cumulative fluorescence in a 0.5s window after each reward or air puff delivery time (+0.05 to +0.55 s) and the cumulative fluorescence in a 0.5s window before each reward or air puff delivery time (−0.5 to 0 s). The response to reward or air puff is defined as the average of this value for all reward or air puff delivery times in a recording. The response to rewards calculated in this manner was used to sort all reward and air puff triggered average heatmaps in Fig. 4D-F and Fig. S1G; S6A, C-D; S7A. Heatmaps for air puff responses (Fig. 4F, S6E, S7B) were sorted by the corresponding reward responses for each recording, with recordings with no rewards being shown at the top (mice not licking for certain recordings result in a higher number of recordings included for air puff than reward analysis). Fig. 4J-K show the location of each recording color-coded based on the reward or air puff response for that recording, calculated in this manner.

P-values for reporting statistical significance for each subtype’s responses to rewards and air puffs (Fig. 4G, S7C) used a non-parametric statistical test (Wilcoxon signed-rank test), with p-values corrected for the number of comparisons conducted (Bonferroni correction). P-values for reporting sensitivity to reward size (Fig. 2I, S7E) were calculated using a non-parametric paired statistical test (Wilcoxon signed-rank test), with Bonferroni correction, between the responses to small and large rewards in the same recording.

SNc recordings (Fig. S7A-D) were analyzed in the same manner as striatal recordings.

### K-means clustering

K-means clustering was run using the MATLAB *kmeans* function for 3 clusters on the values of reward and air puff responses (see previous section for calculation) and the scores along the first two principal components (PC1, PC2) from the PCA analysis on cross-correlations between ΔF/F and acceleration traces (as in Fig. 2K), for all axonal recordings from Calb1, VGlut2 and Anxa1 subtypes where all measures were obtained (mice were running above threshold and received rewards and aversive stimuli, following the same inclusion criteria as previously explained). From the 3 resulting clusters, each subtype was matched to the cluster with the greatest overlap (each cluster was matched to a different subtype), and accuracy (Fig. 5B) was calculated as the percentage of recordings classified within that cluster. Because this k-means clustering was run on a 4-dimensional dataset (reward, air puff, locomotion PC1 score, locomotion PC2 score), Fig. 5A instead shows the combination of PC1 and PC2 scores as an angle, as calculated above.

### Cross-correlation between SNc and striatum ΔF/F traces

All simultaneously recorded pairs of SNc/striatum recordings where both traces had a signal-to-noise ratio above 10 were included, regardless of behavior. Cross-correlations between SNc and striatum ΔF/F traces were calculated using MATLAB’s *crosscorr* function over a 1s time-lag window (100 bins). For the isosbestic control cross-correlation shown in Fig. 6D-E and Fig. S8B-C, we calculated the cross-correlations between SNc-470 and striatum-405 ΔF/F traces and also between SNc-405 and striatum-470 ΔF/F traces, and averaged the resulting cross-correlation traces together. Any pairs of recordings with a peak 405/470 average cross-correlation above 0.12 were excluded.

For plotting in Fig. 6D-E and Fig. S8B-C, 405 and 470 cross-correlations were smoothed over 5 bins (0.05s). Shaded areas in represent the mean ± standard error of the mean (s.e.m.), while accompanying heatmaps show cross-correlations for all recordings. For comparison of peak cross correlations between each subtype and DAT (Fig. 6F, S8D) we used a non-parametric statistical test for two independent populations (Mann-Whitney U test, also called Wilcoxon rank-sum test), with Bonferroni correction (p-values were multiplied by the number of comparisons performed).

### Fiber placement localization

For the representation of recording locations in striatum shown in Fig. 2E, 3A-B, 4J-L and Fig. S5A, 20x magnification images of striatum were acquired on a Keyence slide scanner (BZ-X810) (see Method details, Histology). For the slice in each brain with the clearest fiber track, fiber tracks were marked onto the images. We then identified the closest reference slice for each imaged brain slice (reference slices from the Paxinos Mouse brain atlas), spaced 0.36 mm (bregma +0.86, +0.50, +0.14, -0.22, -0.58, -0.94, -1.34 mm; as shown in schematics in Fig. S5A), and uniformly scaled this reference to approximately match the imaged slice. Recording locations for recordings included in each figure for each mouse were then marked on each slice, measuring depth from brain surface along the fiber track. The fiber tracks and recording locations mapped to these reference slides from all mice for each subtype were combined for Fig. 2E, 3A, 4L, S5A. Circles represent approximate light collection recording area for our 200 µm fibers (∼300 µm in diameter). For compact representation in Fig. 3B and 4J-K, all slices from the body of the striatum (bregma +0.85 to +0.14) or posterior striatum (bregma -0.58 to -1.34) were approximately aligned and combined.

**Fig. S1:**
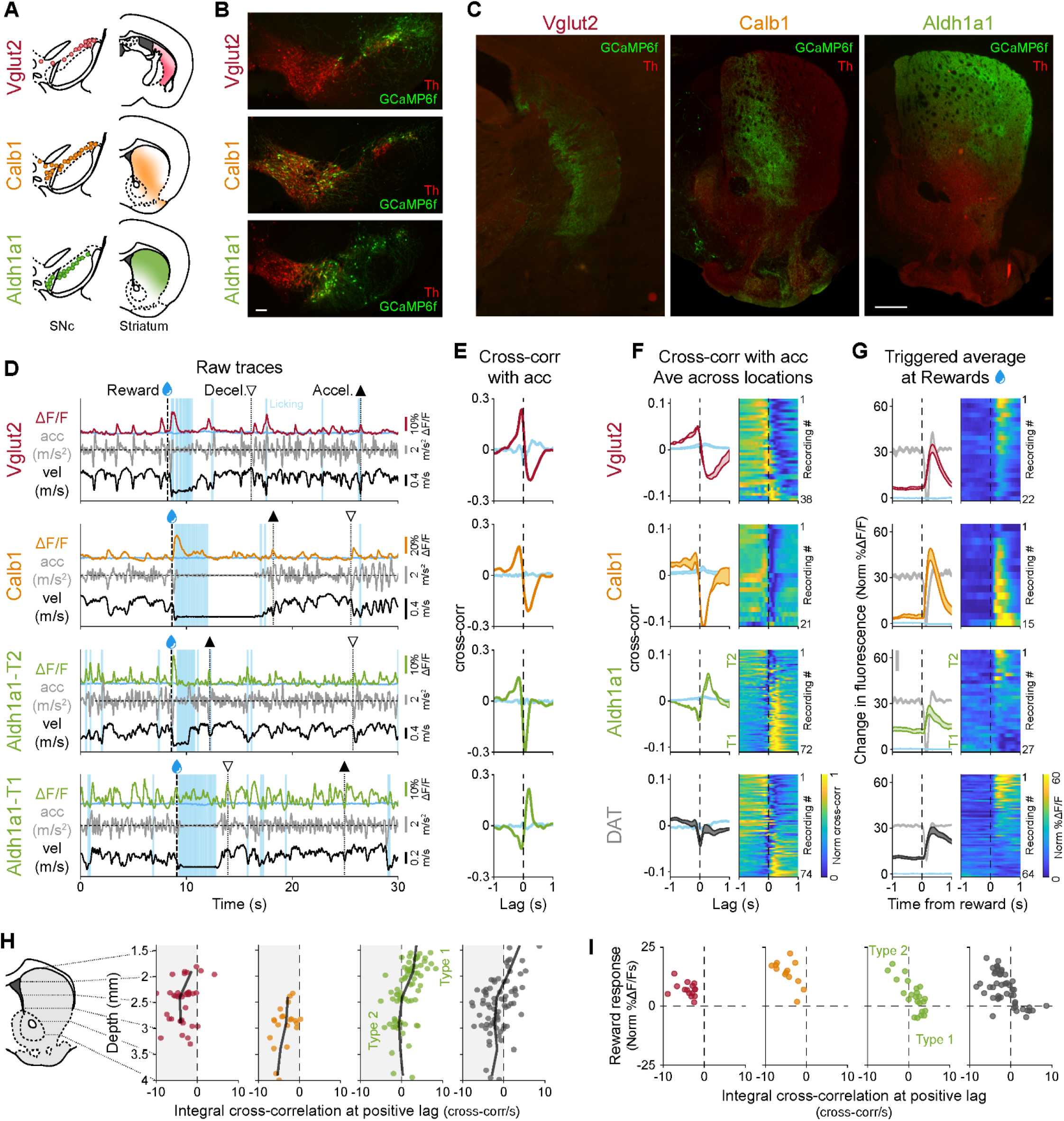
The Aldh1a1+ subtype is functionally heterogeneous. **(A)** Schematic showing the distribution of somas and axons across the SNc and striatum for three previously described subtypes (See Poulin et al. 2018). **(B)** Representative distribution of somas for different subtypes within SNc. Scale bar 100 um. **(C)** Representative projection patterns of different subtypes in striatum. Scale bar 500 um. **(D)** Example recordings for each subtype studied (two from Aldh1a1 with different functional signaling patterns, Type 1 and Type 2), showing fluorescence traces (ΔF/F), velocity, acceleration, licking, and reward delivery times. lsosbestic control shown in blue. Large accelerations= **▴**, large decelerations= Δ. **(E)** Cross-correlation between ΔF/F traces and acceleration for traces shown in D. lsosbestic control shown in blue. **(F)** Average cross-correlation between ΔF/F traces and acceleration for all recordings of each subtype and DAT (subtypes indiscriminately labeled). lsosbestic control shown in blue. Shaded regions denote mean ± s.e.m. Heatmap shows cross-correlations for each recording, sorted by the integral of the cross-correlation at positive lags. Vglut2 mice = 11, n = 38 recordings; Calb1 mice = 5, n = 21 ; Aldh1 a1 mice = 13, n = 72 DAT mice = 14, n = 74. **(G)** ΔF/F triggered averages on reward delivery times for all recordings of each subtype and DAT. lsosbestic control shown in light blue, same scale as ΔF/F average. Acceleration shown in gray in the background (scale bar = 0.2 m/s^2^). Shaded regions denote mean ± s.e.m. Heatmap shows triggered average for each recording, sorted by size of reward response. Vglut2 mice = 10, n = 22 recordings; Calb1 mice = 7, n = 15; Aldh1 a1 mice = 7, n = 27; DAT mice = 12, n = 64. **(H)** Distribution of locmotion response (integral of the cross-correlation at positive lags) along the dorso-ventral axis of the striatum for all recordings of all subtypes and DAT, showing how in Aldh1a1 dorsal recordings show acceleration correlation (Type 1) while more ventral recordings show deceleration correlation (Type 2). Black line represents moving average (0.5 mm bins). **(I)** Relationship between reward response and locomotion response for each recording of each subtype, showing how in Aldh1 a1 larger reward responses correspond with deceleration correlation (Type 2), while small or negative reward responses correspond with acceleration correlation (Type 1).

**Fig. S2:**
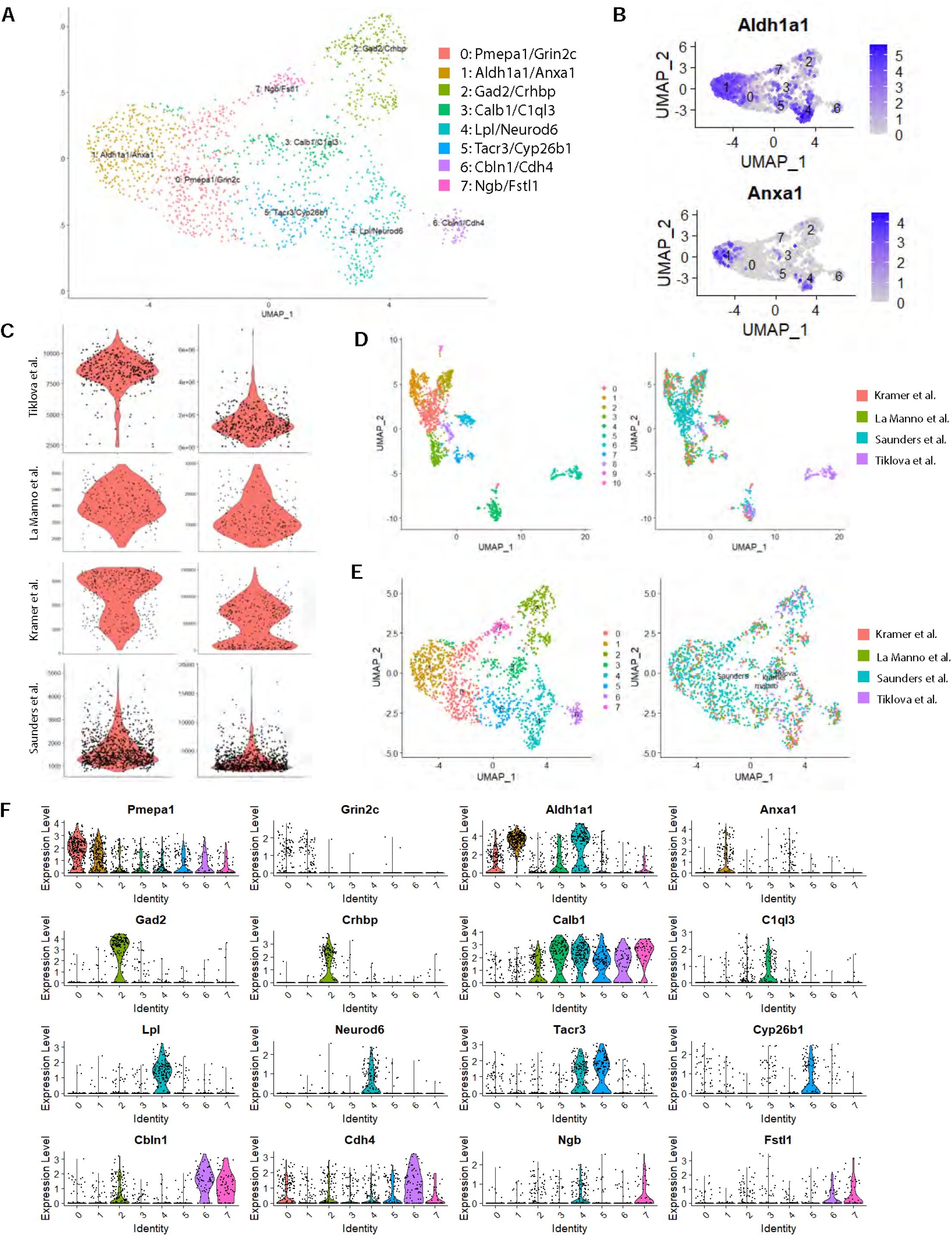
Integration of scRNA-seq datasets reveals more granular resolution of DA neuron subtypes. **(A)** Resulting clusters from integrating datasets. **(B)** Expression patterns of Anxa1 and Aldh1a1, the top defining markers for cluster 1. Expression of Anxa1 appear to be limited to a subset of Aldh1a1-expressing neurons. **(C)** Violin plots of number of genes and RNA counts from each source dataset, which were used to determine cutoffs for quality control filtering. **(D)** LIGER clustering of the meta-dataset, revealing one cluster that was more distantly related to all other DA neurons and came solely from the Tiklova et al. dataset. This cluster was subsequently removed. (E) Cells colored by cluster (left) or source dataset (right), which reveals that all clusters were represented by each dataset. **(F)** Violin plots of the top 2 defining marker genes for each cluster.

**Fig. S3:**
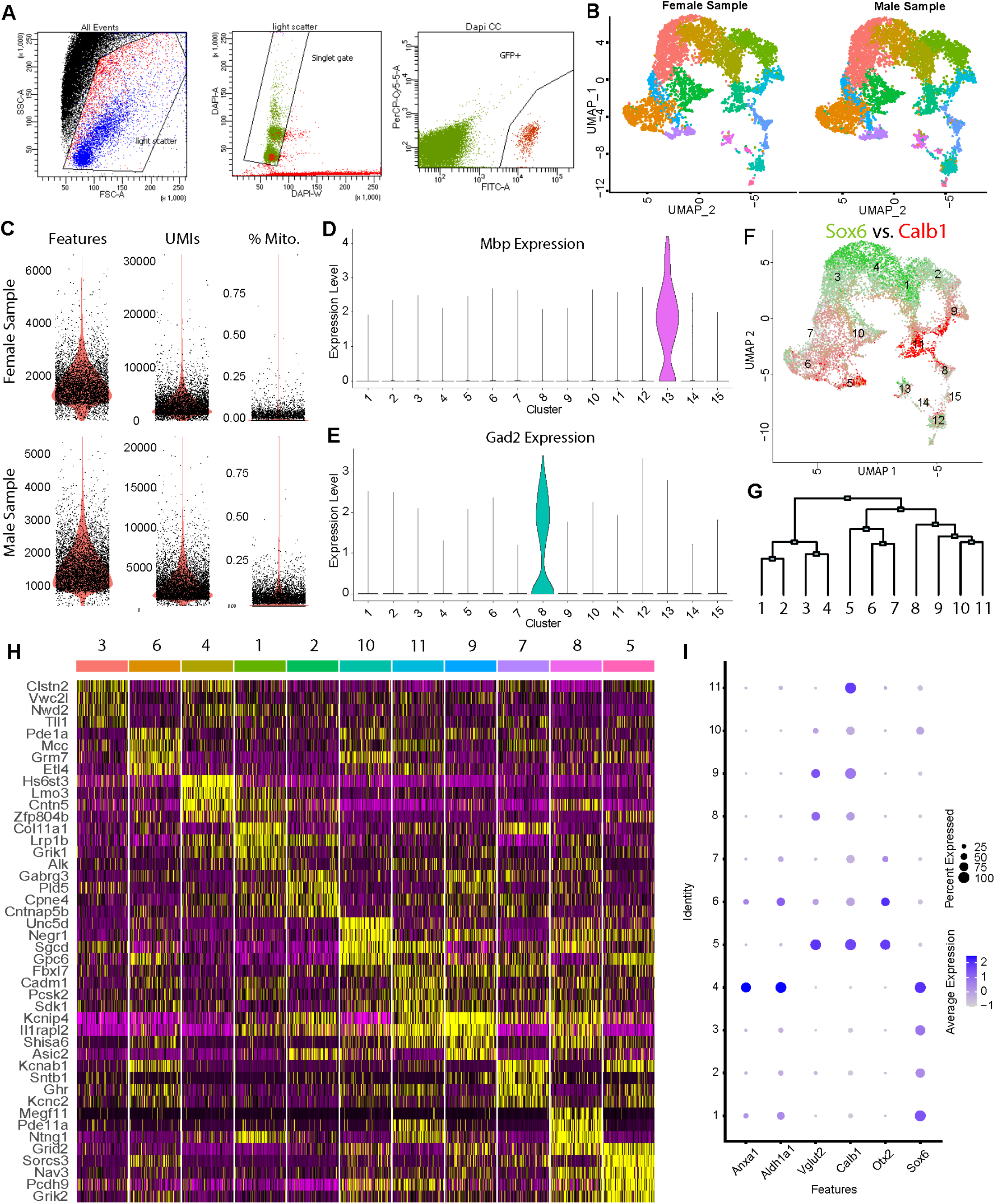
Generation and analysis of single-nucleus RNAseq dataset. **(A)** Example plots from FACS sorting of GFP+ nuclei. **(B)** Plots showing the distribution of cells from either the male or female samples, showing all clusters were represented by both samples. **(C)** Quality control plots of number of genes (features), UMls, and percent mitochondrial reads for each sample. **(D)** Violin plots of Mbp, showing significant expression in cluster 13. **(E)** Expression of Gad2, which is limited to cluster 8, suggesting this cluster represents a previously described population of dopamine neurons with some GABAergic characteristics. **(F)** Overlaid expression patterns of Sox6 (green) and Calb1 (red) recapitulates a previously observed dichotomy among midbrain dopamine neurons. **(G)** Diagram of hierarchical clustering estimation. Notably, clusters 1-4 appear to be Sox6+, 5-7 are Otx2+, and 8-11 are negative for both markers. **(H)** Heatmap of top 4 differentially expressed genes for each cluster, excluding clusters 12, 13, 14 and 15, which do not appear to be classic midbrain dopamine neurons based on lower expression of pan-DA neuron markers (TH, DOC, DAT and Vmat2) (I) Dotplot of expression (post zero-imputation) of several key marker genes of dopamine neuron subpopulations.

**Fig. S4:**
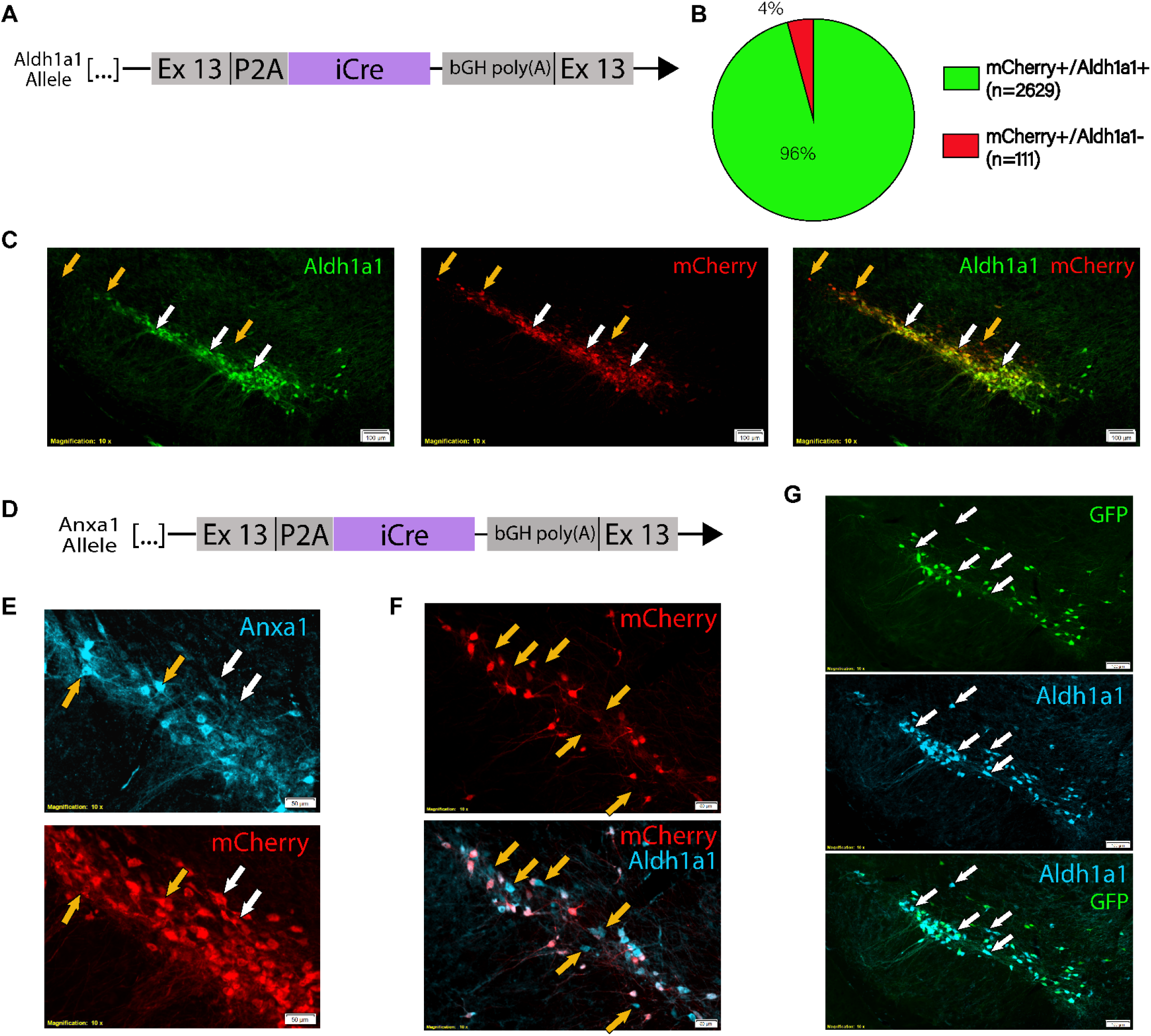
Validation of Aldh1a1-iCre and Anxa1-iCre mouse lines. **(A)** Schematic representation of Aldh1a1-iCre transgenic line. Endogenous Aldh1a1 gene was targeted for insertion of a P2A peptide and iCre immediately following the peptide encoded by Exon 13. **(B)** Ratio of mCherry virally labeled cells co-staining for Aldh1a1 (n=4 mice). (C) Substantia nigra pars compacta immunofluorescence staining from Aldh1a1-iCre mice injected with an AAV5-DIO-mCherry virus. Co-staining shows excellent efficiency and fidelity of iCre recombination, which is notably limited to TH+ cells in this region. White arrows: examples of mCherry and Aldh1a1 co-stained cells. Orange arrows: mCherry-ex pressing cells with undetectable Aldh1a1 staining, which were primarily localized to the dorsal and lateral SNc. **(D)** Schematic representation of Anxa1-iCre transgenic line. **(E)** High magnification of immunofluorescence staining fromAnxa1-iCre mice injected with an AAV5-D10-mCherry virus shows that recombination occurs in cells with both high Anxa1 protein staining (orange arrows) as well as low Anxa1 protein (white arrows). (F) IF staining for Aldh1a1 in the same virally labeled brains as previous panel shows thatAnxa1-iCre mediated recombination occurs in only a subset of Aldh1a1-expressing neurons. Examples of Aldh1a1+/mCherry-cells are shown with yellow arrows. **(G)** IF staining of GFP and Aldh1a1 in Anxa1-iCre, TH-Flpo, RC::FrePe mice. Recombination by iCre and Flpo leads to GFP expression in Anxa1 + DA neurons. Co-staining with Aldh1a1 corroborates that Anxa1-iCre recombination is less broad than Aldh1a1 expression and confirms that viral labeling results were not due to insufficient viral delivery/ diffusion (example of Aldh1a1+, GFP-cells shown with white arrows).

**Fig. S5:**
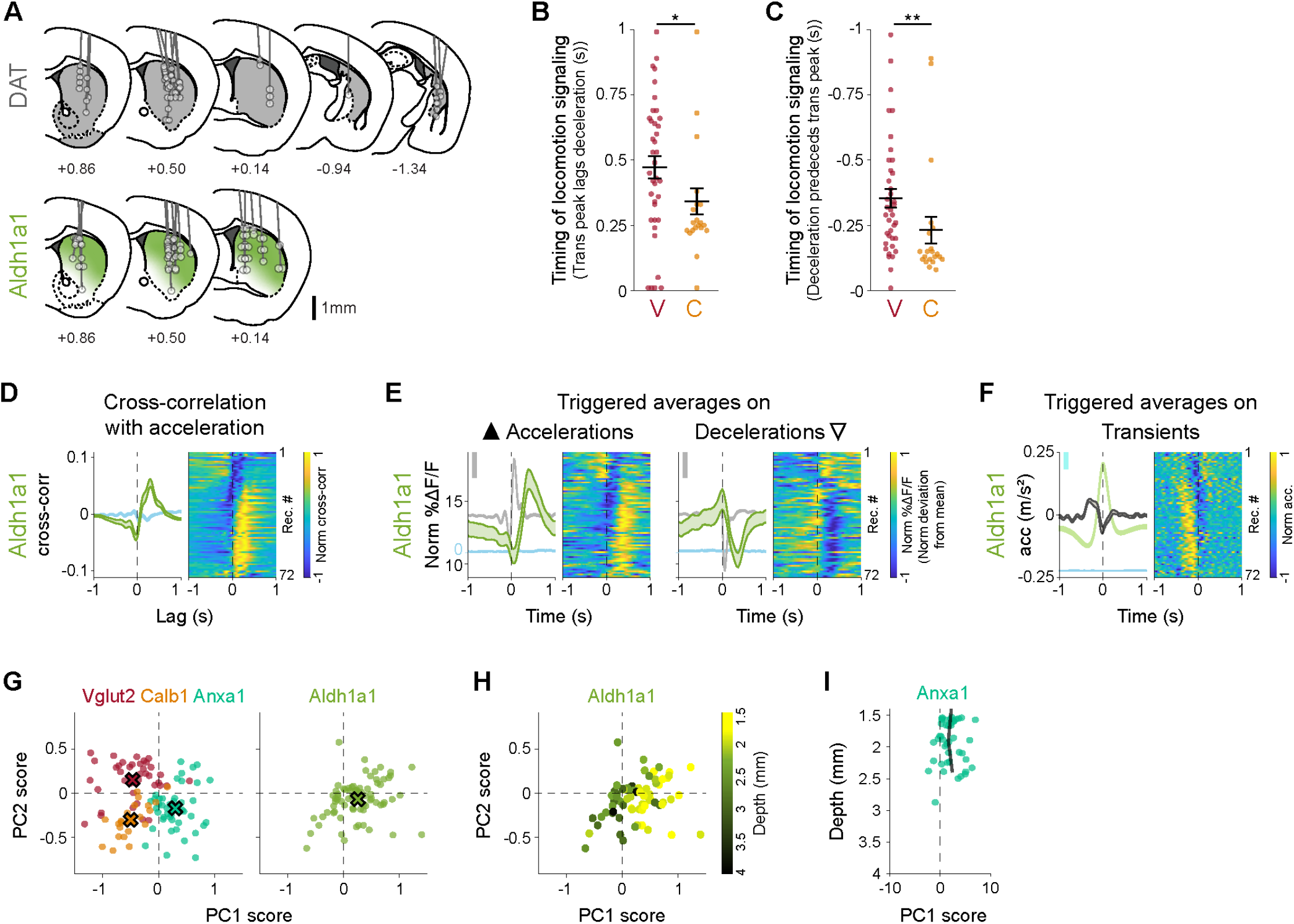
Dopaminergic subtypes show different signaling patterns during locomotion. **(A)** Recording locations for DAT mice and Aldh1a1. **(B)** Timing of the calcium transient peak in triggered averages on decelerations (Fig. 2G, right) for each recording from Calb1 and Vglut2. Means Vglut2 = 0.35, Calb1 = 0.23; p-value for comparison between subtypes= 0.02 (Wilcoxon rank-sum test with Bonferroni correction). **(C)** Timing of the deceleration peak in triggered averages on ΔF/F transient peaks (Fig. 2H) for each recording from Calb1 and Vglut2. Means Vglut2 = 0.47, Calb1 = 0.34; p-value for comparison between subtypes= 0.001 (Wilcoxon rank-sum test with Bonferroni correction). **(D)** Average cross-correlation between ΔF/F traces and acceleration for Aldh1a1 recordings (as Fig. 2F). lsosbestic control shown in blue. Shaded regions denote mean ± s.e.m. Heatmap shows cross-correlation for each recording, sorted by PC1/PC2 angle (see Fig. 2L). Aldh1a1 mice = 13, n = 72 recordings. **(E)** Triggered averages on large accelerations (left, **▴**) and large decelerations (right, ▿) for Aldh1a1 recordings (as Fig. 2G). lsosbestic control shown in light blue, same scale as ΔF/F average but shifted. Acceleration shown in grey in the background (scale bar = 0.2 m/s^2^). Shaded regions denote mean ± s.e.m. Heatmap shows triggered average for each recording, sorted as in D. **(F)** Triggered averages on large transients for Aldh1a1 recordings (as Fig. 2H). ΔF/F average and isosbestic control shown in the background (scale bar = 5% Norm ΔF/F.) Shaded regions denote mean ± s.e.m. Heatmap shows triggered average for each recording, sorted as in D. **(G)** Principal component scores for each recording along PC1 and PC2, comparing Aldh1a1 subtypes (right in green) to the other subtypes as shown in Fig. 2K (left). X shows mean for each subtype. **(H)** Same as G right but each Aldh1a1 recording is color-coded by depth within striatum, showing that Aldh1a1 axons deeper in striatum show similar locomotion signaling to Calb1. **(I)** Distribution of locomotion response (integral of the cross-correlation at positive lags) along the dorso-ventral axis of the striatum forAnxa1 (as Fig. S1H). Black line represents moving average (0.5 mm bins).

**Fig. S6:**
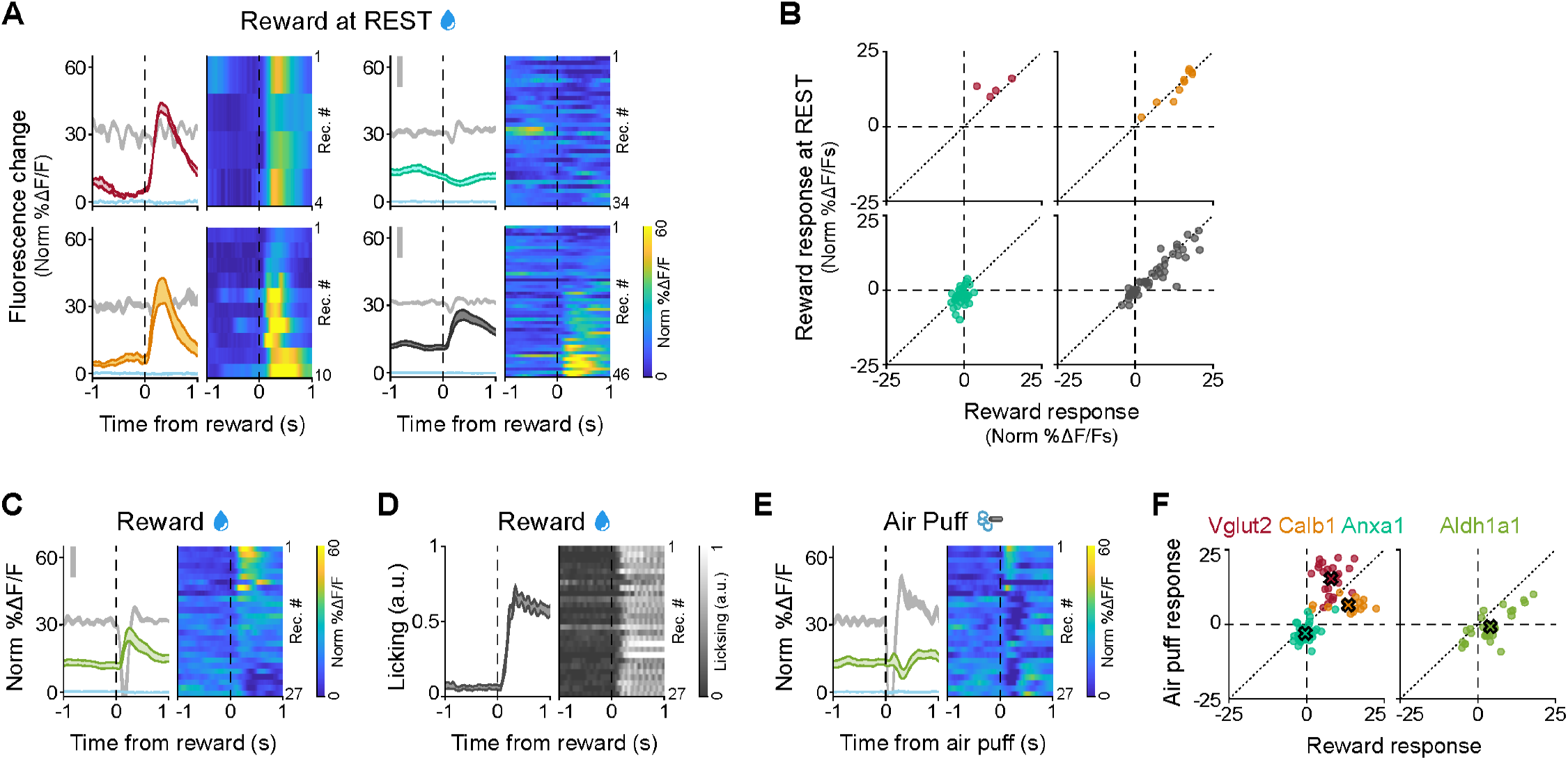
Dopaminergic subtypes show different signaling to rewards and aversive stimuli. **(A)** ΔF/F averages triggered on rewards delivered during rest for all recordings of each subtype and DAT. lsosbestic control shown in light blue, same scale as ΔF/F average. Acceleration shown in gray in the background (scale bar = 0.2m/s^2^). Shaded regions denote mean ± s.e.m. Heatmaps show triggered average for each recording, sorted by size of the reward response. Vglut2 mice = 3, n = 4 recordings; Calb1 mice = 7, n = 10; Anxa1 mice = 6, n = 34; DAT mice = 11, n = 46. **(B)** Comparison between response to rewards at rest (A) vs response to all rewards (Fig. 4D) for all recordings of each subtype and DAT. Diagonal dotted line represents identity line (same response to rewards at rest vs all rewards). **(C)** ΔF/F average triggered reward delivery times for all recordings from Aldh1a1 (as Fig. 4D). lsosbestic control shown in light blue, same scale as ΔF/F average. Acceleration shown in gray in the background (scale bar = 0.2m/s^2^). Shaded regions denote mean ± s.e.m. Heatmap shows triggered average for each recording, sorted by size of reward response. Aldh1a1 mice = 7, n = 27 recordings. **(D)** Licking average triggered on reward delivery times (same as D) for all recording from Aldh1a1 (as Fig. 4E). Shaded regions denote mean ± s.e.m. Heatmap shows triggered average for each recording, sorted as in C. **(E)** ΔF/F average triggered on air puff delivery times for all recordings from Aldh1a1 (as Fig. 4F). lsosbestic control shown in light blue, same scale as ΔF/F average. Acceleration shown in gray in the background (scale bar = 0.2m/s^2^). Shaded regions denote mean ± s.e.m. Heatmap shows triggered average for each recording, sorted by reward size as in C, D. Aldh1a1 mice = 7, n = 27 recordings. **(F)** Reward vs air puff responses comparing Aldh1a1 subtypes (right in green) to the other subtypes shown in Fig. 4H (left). X shows mean for each subtype.

**Fig. S7:**
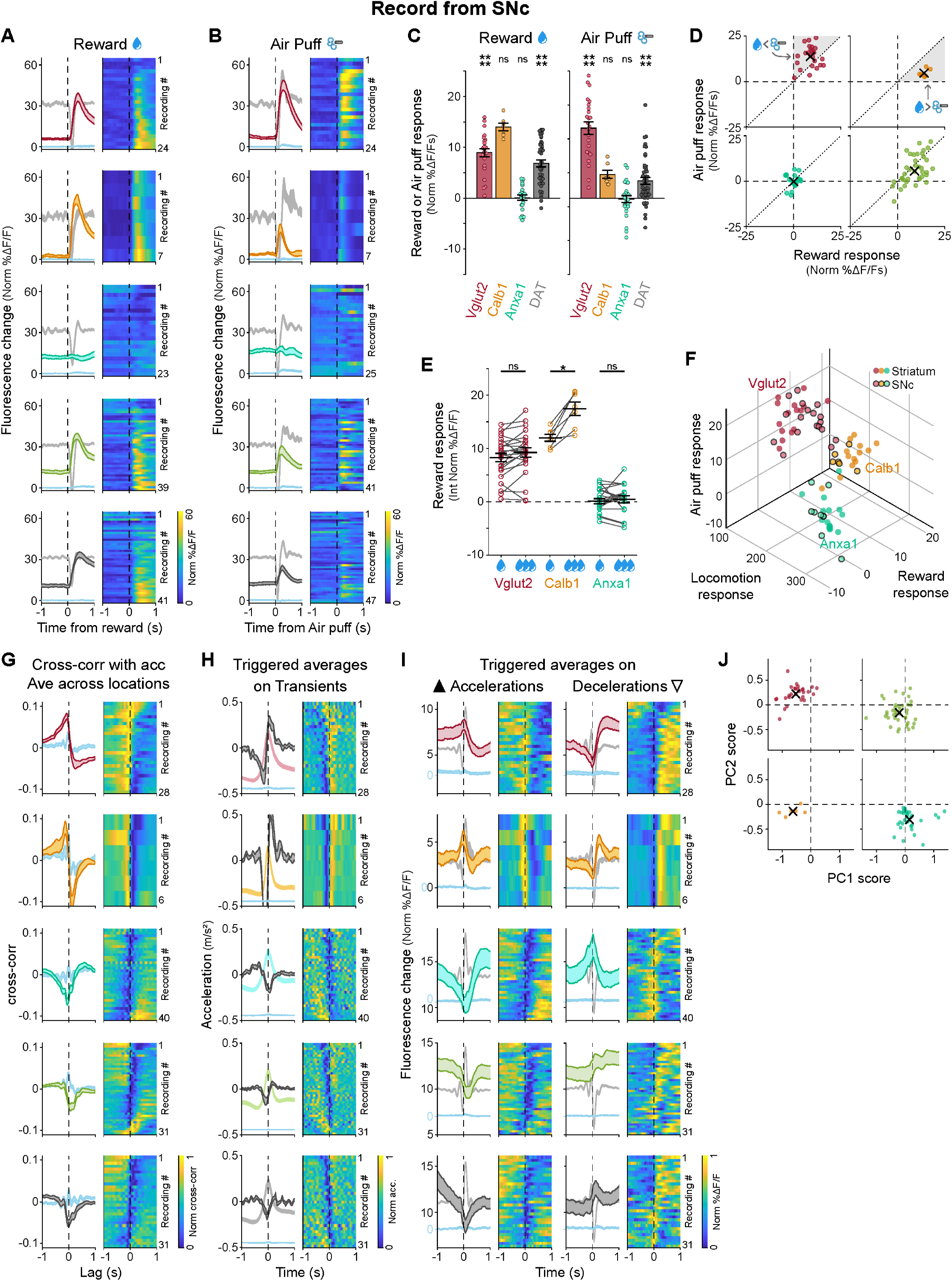
SNc somas of genetic dopamine neuron subtypes have similar signaling patterns to their axons in response to rewards and air puffs and during locomotion. **(A-E)** Same as Fig. 4 but for recordings made in SNc. **(A)** ΔF/F averages triggered on reward delivery limes for all recordings of each subtype and DAT. lsosbestic control shown in light blue, same scale as ΔF/F average.Acceleration shown in gray in the background (scale bar = 0.2 m/s^2^). Shaded regions denote mean ± s.e.m. Heatmap shows triggered average for each recording, sorted by size ofreward response. Vglut2 mice = 8, n = 24 recordings; Calb1 mice = 4, n = 7;Anxa1 mice = 6, n = 23; Aldh1a1 mice = 10, n = 39; DAT mice = 8, n = 41. **(B)** ΔF/F averages triggered on air puff delivery limes for all recordings of each subtype and DAT. lsosbestic control shown in light blue, same scale as ΔF/F average.Acceleration shown in gray in the background (scale bar = 0.2 m/s^2^). Shaded regions denote mean ± s.e.m. Heatmap shows triggered average for each recording, sorted by reward size as in A. Vglut2 mice = 8, n = 24 recordings; Calb1 mice = 4, n = 7; Anxa1 mice = 6, n = 25; Aldh1a1 mice = 10, n = 41; DAT mice = 8, n =41. **(C)** Average reward and air puff responses for each subtype. Error bars denote± s.e.m. p-values for reward: Vglut2 = 7 × 10-^05^, Calb1 = 0.06 (not significant), Anxa1 = 1 (not significant), DAT= 5 × 10-^07^.p-values for air puff: VGlut2 = 7 × 10-0s, Calb1 = 0.06, Anxa1 = 1 (not significant), DAT= 3 × 10-^05^. Wilcoxon signed-rank test with Bonferroni correction. **(D)** Reward vs air puff responses for all recordings of each subtype and DAT. X shows mean for each subtype. Shaded regions are areas representing greater air puff than reward response (for Vglut2) or greater reward vs air puff response (for Calb1). **(E)** Comparison ofresponses to small vs large rewards for each subtype. Error bars denote mean ± s.e.m. p-values: Vglut2 = 0.1 (not significant), Calb1 = 0.047, Anxa1 = 1 (not significant). Paired Wilcoxon Signed Rank test with Bonferroni correction. **(F)** 3D plot showing locomotion (PC1/PC2 angle), reward and air puff responses for each recording and each subtype, comparing slriatal recordings (same as Fig. 5A) and SNc recordings. **(G-J)** Same as Fig. 2 but for recordings made in SNc. **(G)** Average cross-correlation between ΔF/F traces and acceleration for all recordings of each subtype. Isosbestic control shown in blue. Shaded regions denote mean ± s.e.m. Heatmap shows corss-correlation for each recording, sorted by PC1/PC2 angle (see Fig. 2L). Vglut2 mice = 11, n = 28 recordings; Calb1 mice = 3, n = 6;Anxa1 mice = 8, n = 34;Aldh1a1 mice = 12, n =41; DAT mice = 8, n = 31. **(H)** ΔF/F averages triggered on large accelerations (left, **▴**) and large decelerations (right, ▿) for all recordings of each subtype. lsosbestic control shown in light blue, same scale as ΔF/F average but shifted. Acceleration shown in gray in the background (scale bar = 0.2 m/s^2^). Shaded regions denote mean ± s.e.m. Heatmap shows triggered average for each recording, sorted as in G. **(I)** Acceleration averages triggered on large transients for all recordings of each subtype. ΔF/F average and isosbestic control shown in the background (scale bar = 5% Norm ΔF/F.) Shaded regions denote mean ± s.e.m. Heatmap shows triggered average for each recording, sorted as in G. **(J)** Principal component scores for each recording of each subtype along PC1 and PC2 (same PCs obtained from the slriatal recordings, as shown in Fig. 2J-L). X shows mean for each subtype.

**Fig. S8:**
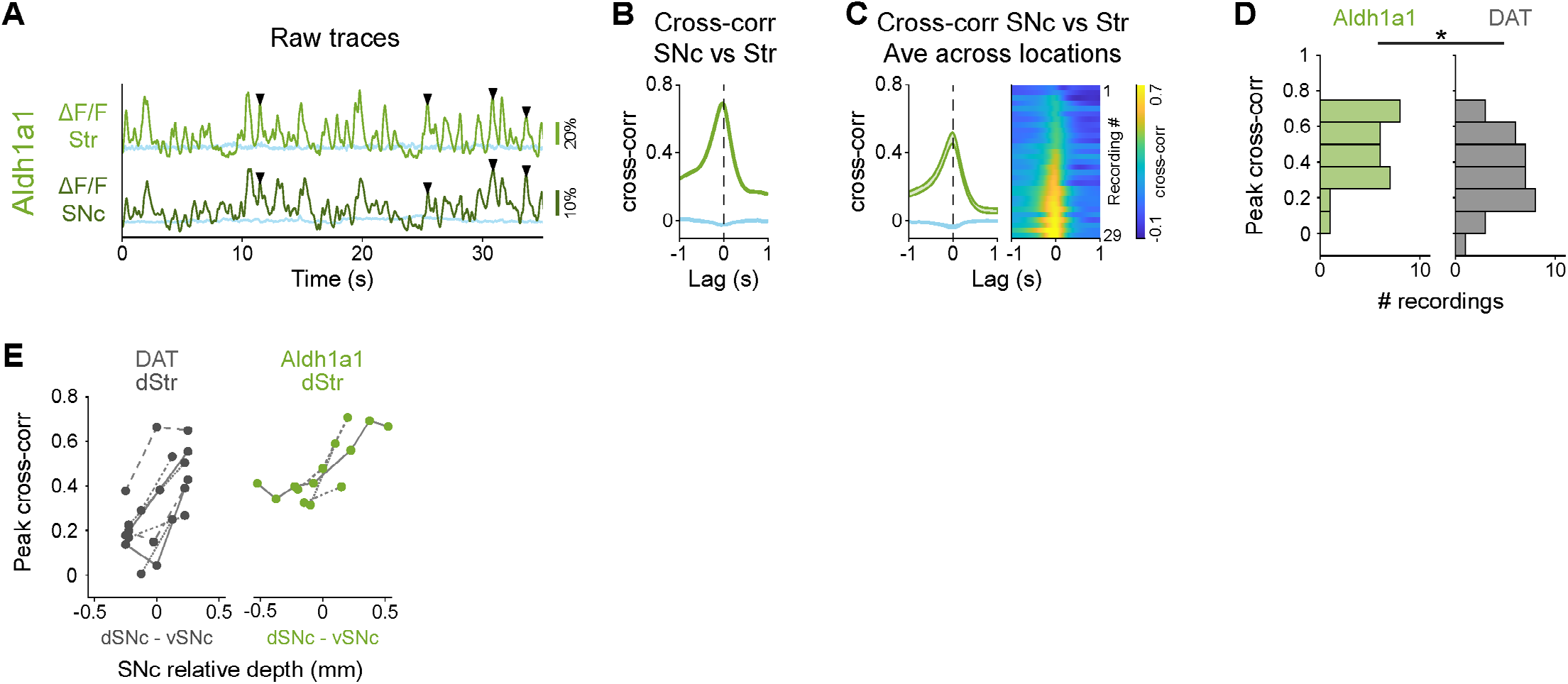
Highly correlated signaling in axons and somas within genetic subtypes of dopamine neurons. **(A)** Example recording for Aldh1a1 showing simultaneous fluorescence traces (ΔF/F) from SNc and striatum. lsosbestic control shown in blue. **▾** = Example transients present in SNc and in striatum. **(B)** Cross-correlation between ΔF/F traces from striatum and SNc shown in A. lsosbestic control shown in blue. **(C)** Average cross-correlation between ΔF/F traces from striatum and SNc for all recordings of Aldh1a1 (as Fig. 6E). lsosbestic control shown in blue. Shaded regions denote mean ± s.e.m. Heatmap shows cross correlations for each paired recording sorted by peak magnitude. Aldh1a1 mice = 8, n = 29 recordings. **(D)** Distribution of peak cross correlations between SNc and striatum for recordings of Aldh1a1 and DAT shown in C (as Fig. 6F). P-value for comparison DAT-Aldh1a1 = 0.03 (Mann-Whitney U test with Bonferroni correction). **(E)** Peak cross correlations between dorsal striatum recordings (from Aldh1a1 or DAT) vs different relative depths in SNc, showing that for Aldh1a1 dorsal striatum signaling is best correlated to ventral SNc.

